# Two closely related β-1,2-xylosyltransferases differentially impact cryptococcal glycan synthesis

**DOI:** 10.64898/2026.04.21.719979

**Authors:** Daphne Boodwa-Ko, J. Stacey Klutts, Kazuhiro Aoki, Michael L. Skowyra, Indrani Bose, Daniel P. Agustinho, Mayumi Ishihara-Aoki, Michael Tiemeyer, Tamara L. Doering

## Abstract

*Cryptococcus neoformans* is an opportunistic fungal pathogen that causes pulmonary infection in immunocompromised patients, which in severe cases leads to fatal meningoencephalitis. *Cryptococcus* exhibits unique glycobiology that plays important roles in pathogenesis. Unlike model yeast and other common fungal pathogens, *Cryptococcus* incorporates xylose, a five-carbon monosaccharide, into its glycans. One trimer motif, which consists of xylose in β-1,2 linkage to the reducing mannose of an α-1,3-mannose dimer, occurs in key cryptococcal glycoconjugates that include protein *N*- and *O*-linked glycans, glycosylinositol phosphorylceramides (GIPCs), and the capsule polysaccharides glucuronoxylomannan (GXM) and glucuronoxylomannogalactan (GXMGal). We previously identified <u>c</u>ryptococcal β-1,2-<u>x</u>ylosyltransferase 1 (Cxt1), which catalyzes formation of this motif in GIPCs, GXM, and GXMGal. Here, we report the discovery of a second enzyme, <u>c</u>ryptococcal β-1,2-<u>x</u>ylosyltransferase 2 (Cxt2). Through characterization of cells that lack one or both corresponding genes (*CXT1* and *CXT2*), we have dissected the biological roles of these enzymes, which are overlapping but not identical. Notably, Cxt1 and Cxt2 colocalize in the Golgi, influence capsule in a strain-dependent manner, and together are responsible for all xylose addition to *O*-glycans. Overall, our work highlights unique roles of these two enzymes and fills a gap in understanding of cryptococcal glycan synthesis.

**IMPORTANCE:** *Cryptococcus neoformans* is an opportunistic fungal pathogen that causes almost 150,000 deaths each year worldwide. *Cryptococcus* synthesizes unique glycan structures that play important roles in its biology and pathogenesis. One abundant component of these structures is xylose, a five-carbon monosaccharide. Because xylose is not used by many fungal organisms, including model yeast, we have limited information about how cells add it to their glycans. Here we report a xylosyltransferase enzyme that performs this function, and we characterize specific biological roles of this protein and a closely related one we discovered earlier. We find that these proteins together perform all detectable xylose addition to an important class of protein-linked glycans (*O*-glycans). They also both participate in other synthetic processes, although this varies with the specific enzyme and strain background. These results contribute to our understanding of cryptococcal glycan synthesis and underscore the importance of testing multiple background strains.

## INTRODUCTION

*Cryptococcus neoformans* is an opportunistic fungal pathogen found ubiquitously in the environment. This haploid yeast primarily infects immunocompromised patients, such as HIV/AIDS patients and transplant recipients (1). *C. neoformans* first enters the lungs as spores or desiccated yeast and establishes a pulmonary infection. It can subsequently disseminate to the central nervous system, where it causes cryptococcal meningoencephalitis. There are nearly 200,000 cases of cryptococcal meningoencephalitis annually, leading to approximately 150,000 deaths (1). Current treatment options for this devastating disease are limited (2), although new therapies are being explored.

*C. neoformans* is characterized by a polysaccharide capsule which surrounds its fungal cell wall. The capsule consists of two large polysaccharides: glucuronoxylomannan (GXM) and glucuronoxylomannogalactan (GXMGal) (3). GXM, which makes up ∼90% of capsule mass, consists of an α-1,3-mannan backbone with variable glucuronic acid and xylose substitutions (Fig. S1A). GXMGal is comprised of an α-1,6-galactan backbone bearing galactomannan sidechains that are substituted with glucuronic acid and xylose (Fig. S1B) (4, 5). The capsule markedly expands when *Cryptococcus* enters a mammalian host or is grown *in vitro* under conditions that mimic the host environment (6–8). Capsule polysaccharides are essential for virulence and play important roles in modulating the host immune response (9–15).

*C. neoformans* has historically been categorized into four serotypes (A-D) (3, 4, 16–18). Serotypes B and C correspond to *C. gattii* and *C. bascillisporus*, respectively, while serotypes A and D correspond to two varieties of *C. neoformans*, *var. grubii* and *var. neoformans.* These have also been named *C. neoformans* and *C. deneoformans* (19), the nomenclature we use below. *C. neoformans* is the most common *Cryptococcus* species in environmental, clinical, and veterinary isolates and makes up 63% of all collected isolates. *C. deneoformans* is much less common, accounting for 4-5% of isolates (19). This disparity may reflect differences in cellular properties and virulence, with *C. neoformans* generally more thermotolerant and more virulent than *C. deneoformans* (19, 20). Their capsule GXM composition also differs: *C. neoformans* GXM typically exhibits more xylose substitutions than *C. deneoformans* GXM (4). Despite these distinctions, the two species are closely related, with near complete synteny and 85-90% nucleotide identity (21). Both have been used extensively in cryptococcal research.

Apart from the capsule, *C. neoformans* synthesizes multiple other glycoconjugates, including glycolipids, glycoproteins, and cell wall glycans. Generating these structures requires the synthesis and transport of nucleotide sugars, which act as donors for many glycan biosynthetic processes (22). The transfer of saccharide moieties from nucleotide sugar donors to specific acceptor molecules is catalyzed by glycosyltransferases (GTs). These critical enzymes are generally specific for the donor and acceptor molecules, and form specific glycosidic linkages. Importantly, while amino acid sequences can be predicted to encode GTs, their specific enzymatic activity cannot be accurately determined without biochemical assays. Despite significant research on cryptococcal GTs (23, 24), we do not yet know the full array of enzymes that build cryptococcal glycoconjugates.

Xylose is a five-carbon monosaccharide that does not occur in the model yeast *Saccharomyces cerevisiae* (25, 26), although it has been reported in *C. neoformans* and a handful of other fungal species: the basidiomycetes *Trichosporon asahii* and *mucoides* and the ascomycete *Paracoccidioides brasiliensis* (27–30). It is notably abundant in cryptococcal capsule polysaccharides, *N*- and *O*-linked glycoproteins, and glycosylinositol phosphorylceramides (GIPCs). In many of these, xylose occurs in a trimer motif linked β-1,2 to α-1,3-linked mannose, including in capsule polysaccharides, *N*- and *O*-linked glycoproteins, and glycosphingolipids (Fig. S1, bold text) (26–30).

In *Cryptococcus*, xylose modifications play significant roles in virulence, immune modulation, and cell surface integrity (32, 36–39). The absence of proteins that participate in the synthesis or transport of UDP-xylose, the specific donor for xylosyltransferase enzymes, leads to altered capsule structure and dramatically reduced virulence (32, 36). Conversely, mutants with increased proportions of xylose-containing *N*- and *O*-linked glycans have enhanced immunogenicity compared to their non-xylosylated counterparts (38, 39). Defining the enzymes which create these structures is thus important for understanding this pathogen.

In search of capsule synthetic enzymes, we previously identified a cryptococcal β-1,2-xylosyltransferase, Cxt1. This adds xylose in β-1,2 linkage to the reducing mannose residue of α-1,3-mannobiose *in vitro* (40), forming the common triad noted above. Deletion of the corresponding gene (*CXT1*) eliminates GIPC xylosylation and yields reduced xylose in GXM and GXMGal (33, 41), but does not fully eliminate capsule xylosylation. This suggests the existence of at least one other xylosyltransferase enzyme with similar activity. Here we report on such an enzyme, which we have named Cxt2, and use biochemical, phenotypic, and imaging approaches to define its role in cryptococcal biology. We also demonstrate the role of both enzymes in *O*-glycan synthesis.

## RESULTS

### *C. neoformans* expresses a second β-1,2-xylosyltransferase

We previously identified Cxt1 as a cryptococcal β-1,2-xylosyltransferase responsible for adding xylose to capsule polysaccharides and glycosylinositol phosphorylceramides (GIPCs) (33, 40, 41). *In vitro*, Cxt1 catalyzes the transfer of xylose from UDP-xylose to the reducing mannose of an α-1,3-mannobiose (Man_2_) acceptor, forming a β-1,2 linkage (Fig. 1A). This product is readily observed in assays with wild-type strains (Fig. 1B, left panel). Interestingly, we still observed a small amount of this product in assays of membranes from *cxt1*Δ deletion strains (Fig. 1B, right). We speculated that its synthesis was catalyzed by another *C. neoformans* enzyme, which we wished to identify.

**FIG. 1.**
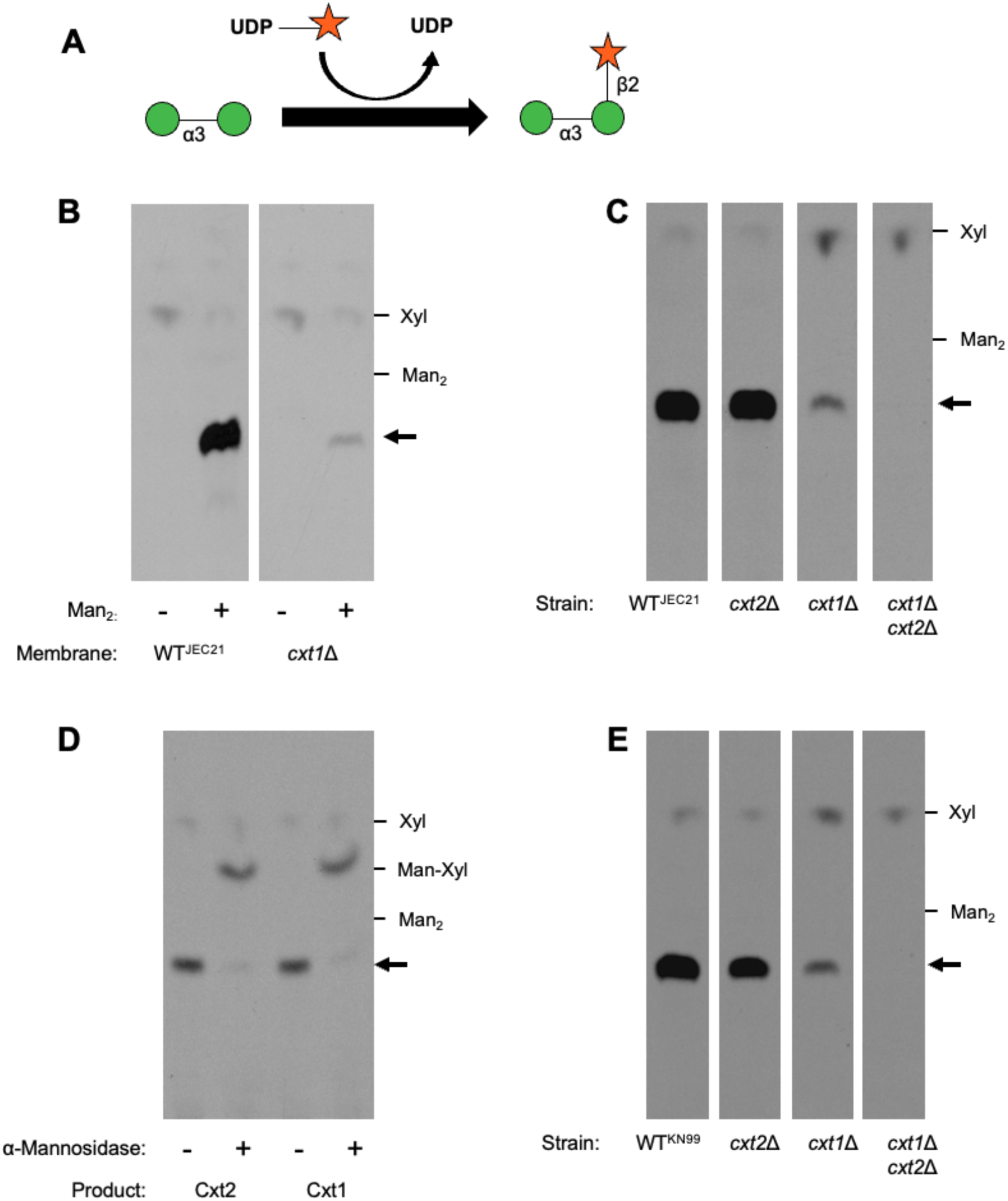
β-1,2-xylosyltransferase activities. (A) Schematic of the β-1,2-xylosyltransferase activity assay. Cxt1 catalyzes the transfer of xylose from UDP-xylose to α-1,3-mannobiose (Man_2_), forming a β-1,2-linkage as shown. Green circle, mannose; orange star, xylose. (B-E) Autoradiographs of assay products resolved by TLC. Migration positions of non-radiolabeled standards are indicated at the right; the black arrow indicates the migration position of the Cxt1 product. WT, wild-type strain; superscript indicates strain background, either JEC21 (*C. deneoformans*) or KN99 (*C. neoformans*). (B-C, E) Washed membranes from the indicated strains were assayed as described in the Methods. (D) Reaction products from assays as in panel (C) were mock treated or incubated with Jack Bean α-mannosidase (see Methods) before TLC.

One candidate for a potential second β-1,2-xylosyltransferase was *CAP5* (gene ID: CND04030), the closest homolog of *CXT1* in the *C. neoformans* genome with 40% amino acid identity and 56% similarity (40, 42). Targeting this gene by RNAi in a *cxt1*Δ background eliminated the residual xylosyltransferase activity, while targeting another homolog, *CAP4* (gene ID: CNB02330), did not (Fig. S2). We hypothesized that the *CAP5* sequence encoded an additional β-1,2-xylosyltransferase.

To test this hypothesis, we deleted the *CAP5* coding sequence, generated isogenic wild-type and mutant strains by mating, and assessed β-1,2-xylosyltransferase activity. While deletion of this gene alone (strain labeled *cxt2*Δ) had little or no effect on the amount of product formed (Fig. 1C, compare first two lanes), its deletion in the *cxt1*Δ background eliminated the residual assay product previously observed in that mutant (Fig. 1C, compare last two lanes). Based on these results, which were consistent with our RNAi experiments, we concluded that this sequence encoded a second β-1,2-xylosyltransferase and renamed it *CXT2*, for <u>c</u>ryptococcal <u>x</u>ylosyltransferase <u>2</u>.

Our prior analysis of the Cxt1 reaction product identified it as α-1,3-mannobiose with β-1,2-xylose on the reducing mannose (40). Because the low abundance of the Cxt2 product (Fig. 1C) made it challenging to isolate sufficient material for such analysis, we instead compared chromatographic behavior of the two assay products. Their comigration on a sensitive TLC system, which resolves multimers of different sizes as well as of different linkages (43, 44), already suggested they had the same structure. To complement these studies, we isolated the Cxt1 and Cxt2 reaction products from *cxt2*Δ and *cxt1*Δ strains respectively, and treated them with Jack Bean mannosidase, which cleaves α-mannose linkages including α-1,3. Each of the resulting digestion products, which we know for Cxt1 is Xyl-β-1,2-Man (40), also comigrated (Fig. 1D). Together, these results show that both Cxt1 and Cxt2 are *C. neoformans* β-1,2-xylosyltransferases.

### Cxt1 and Cxt2 are well-conserved between *C. neoformans* serotypes

Although we performed our early experiments in the *C. deneoformans* strain JEC21, we were interested in whether Cxt1 and Cxt2 were conserved in *C. neoformans*, which causes most human disease and is more virulent. Sequence alignments showed that Cxt1 from JEC21 and *C. neoformans* strain KN99 (45) are 93.23% identical and 96.11% similar; Cxt2 is 88.26% identical and 92.18% similar (Fig. S3A-B). We also assessed structural similarity by calculating the template modeling (TM) score based on the AlphaFold3 predicted structures of each protein (Fig. S3C-D). The TM score for JEC21 and KN99 Cxt1 was 0.95, suggesting high structural similarity. The Cxt2 score was lower (0.80), but still well within the range for structurally similar proteins (0.5-1.00) (46).

When we assayed deletion mutants engineered in KN99, we observed the same product and pattern of β-1,2-xylosyltransferase activity as we had in the *C. deneoformans* strain (Fig. 1E). We conclude that the Cxt enzymes make the same product in both species, with Cxt1 producing the majority of the xylosylated product *in vitro* while Cxt2 produces a smaller fraction.

### β-1,2-xylosyltransferases impact temperature sensitivity in JEC21

To better understand the biological roles of Cxt1 and Cxt2, we compared the growth of single and double deletion mutants to wild-type cells in various stress conditions. The JEC21 *cxt1*Δ *cxt2*Δ strain showed increased sensitivity to 37°C and could not grow at this temperature in the presence of 0.01% SDS, a membrane stressor (Fig. 2). The temperature sensitivity was not exacerbated by high salt, which induces osmotic stress, or calcofluor white (CFW), a cell wall stressor (Fig. S4). The single mutants in this background showed a mixture of phenotypes under stress. Compared to wild type, *cxt1*Δ grew more robustly at 37°C but was more sensitive to SDS. The *cxt2*Δ mutant resembled wild type in most conditions, although it, like *cxt1*Δ, grew worse than wild type in the combined stress of SDS and 37°C. It also grew slightly less well than wild type in DMSO and CFW at that temperature (Fig. S4). In contrast, all KN99 strains grew comparably under all these conditions. Since *C. neoformans* strains are more temperature resistant than *C. deneoformans*, we performed these experiments at 39°C. We observed no difference in growth among KN99 strains (data not shown).

**FIG. 2.**
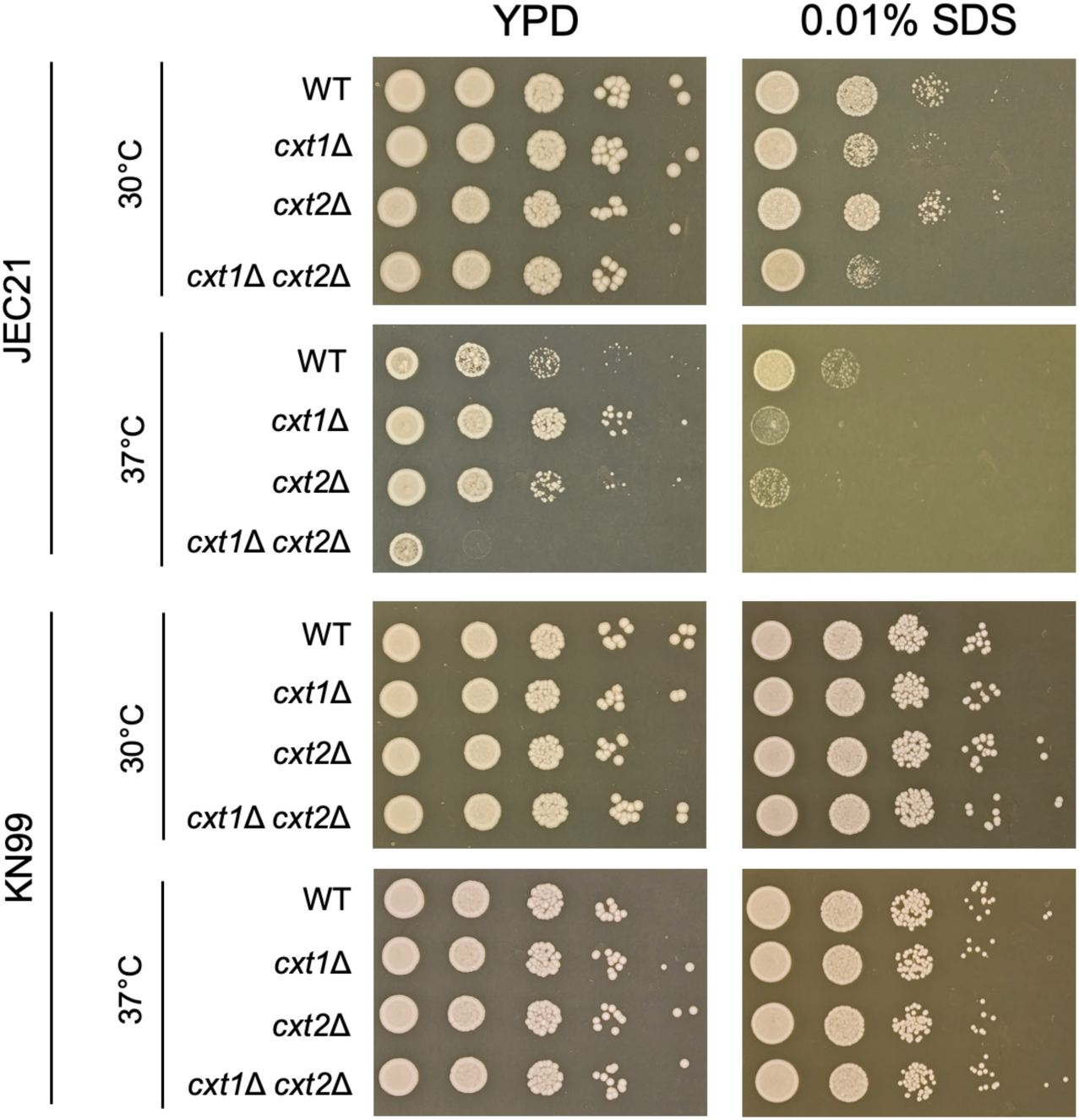
Impact of β-1,2-xylosyltransferases on growth *in vitro*. The indicated strains were plated in 10-fold serial dilutions on YPD agar medium −/+ 0.01% SDS and incubated for 72 h as shown. The experiment shown is representative of three similar studies.

### Cxt1 but not Cxt2 impacts glycosphingolipid synthesis

The SDS sensitivity of the JEC21 mutants suggested a defect in cell membranes, consistent with our earlier finding that Cxt1 catalyzes the addition of xylose to glycosylinositol phosphorylceramides (GIPCs) (33). To assess any effect of Cxt2 on this process, we isolated GIPCs from our JEC21 strain set, as well as from a *uxs1*Δ control strain that cannot form UDP-xylose and therefore lacks all glycosphingolipid xylosylation (31). High-performance TLC (HPTLC) of these samples confirmed that the GIPC profile of *cxt1*Δ cells matches that of the *uxs1*Δ control, indicating they contain no xylose (Fig. S5; note loss of xylosylated-GIPC and accumulation of non-xylosylated-GIPC); results for the double mutant *cxt1*Δ *cxt2*Δ were similar. In contrast, *cxt2*Δ phenocopies wild type, suggesting that Cxt2 has no role in this process. These results are consistent with previous findings that Cxt1 is solely responsible for GIPC xylosylation. However, it also suggests some other reason for the sensitivity of *cxt2*Δ to the combination of SDS and 37°C.

### Cxt1 and Cxt2 add xylose to *O*-linked, but not *N*-linked, glycans

Some *Cryptococcus* mutants defective in *N*- and *O*-linked glycosylation are sensitive to SDS and temperature (34, 35, 38, 47) and studies of *N*- and *O*-glycans in *C. neoformans* suggest that both contain xylose (34, 35, 48). We therefore wondered whether some of the altered growth of our *cxt* mutants might be explained by defects in protein glycosylation.

*N*-linked glycans consist of a branched mannose structure linked to asparagine via a chitobiose core of two *N*-acetylglucosamine residues. In *C. neoformans*, xylose occurs in β-1,2 linkage on the first mannose beyond the core (Fig. S1D). While Cxt1 has been reported to have no direct role in the xylosylation of *N*-glycans (34), Cxt2’s role in this process is not known. When we analyzed *N*-linked glycans in our KN99 strain set by mass spectrometry, the overall profiles were not significantly altered, including the overall abundance of xylose modification (Fig. 3). These results suggest an overall minor impact of Cxt1 and Cxt2 on *N*-glycan xylose modification, consistent with earlier observations on Cxt1 (34). However, compared to wild type or either single mutant, the *cxt1*Δ *cxt2*Δ mutant did exhibit a preponderance of large, xylosylated *N*-glycans (Fig. S6, compare A and B), along with a relative increase in xylosylated N-glycans overall (Fig. S6, panel C). This suggests an indirect effect on *N*-linked glycan biosynthesis, potentially related to donor substrate availability (see Discussion).

**FIG. 3.**
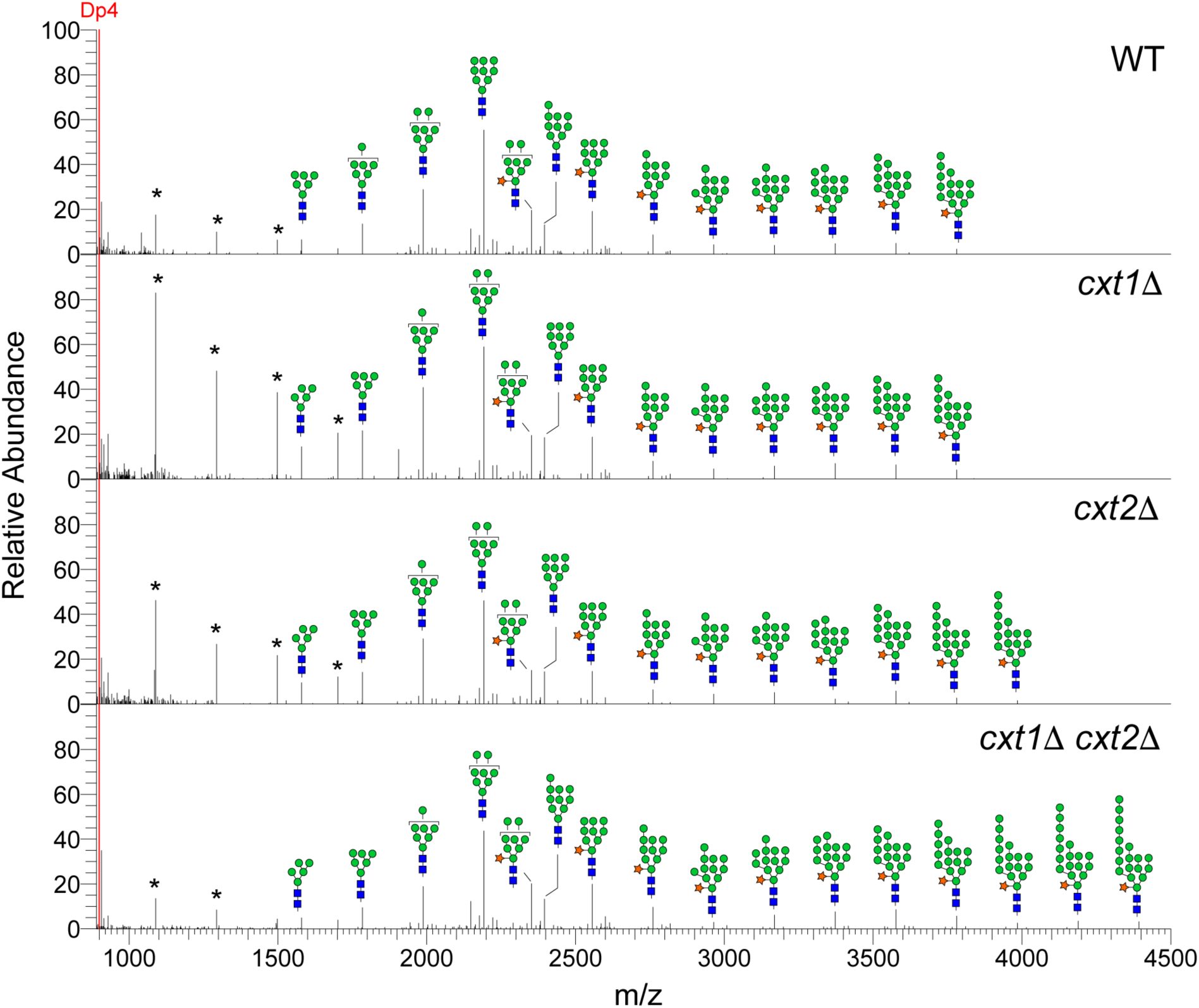
Mass spectrometry analysis of *N*-linked glycans from the indicated strains. DP4 (10 pmol), a maltotetraose standard, was spiked into the *N*-glycan samples for direct comparison. Glycan structures are shown above the corresponding peak; specific m/z values and quantitation are in Table S1. *, common hexose ladder contaminants; blue square, *N*-acetylglucosamine; green circle, mannose; orange star, xylose.

*O*-linked glycans of *C. neoformans* consist of a linear chain of α-linked mannoses, which may be modified with xylose on the reducing terminal mannose residue (Fig. S1E) (35, 48). To assess the roles of Cxt1 and Cxt2 in the synthesis of these structures, we analyzed *O*-glycans isolated from wild type, *cxt1*Δ, *cxt2*Δ, and *cxt1*Δ *cxt2*Δ. Notably, although the effect was greater in *cxt1*Δ, both single mutants showed a significant reduction in xylose-containing structures, including Xyl-Man_2_, Xyl-Man_3_, and Xyl-Man_4_ (Fig. 4A, peaks at 653.36, 857.47, and 1061.58 m/z, respectively; quantified in Fig. 4B). The double mutant showed complete loss of these structures (Fig. 4B). We conclude that, together, Cxt1 and Cxt2 mediate all xylose modification of *O*-linked glycans in *C. neoformans*.

**FIG. 4.**
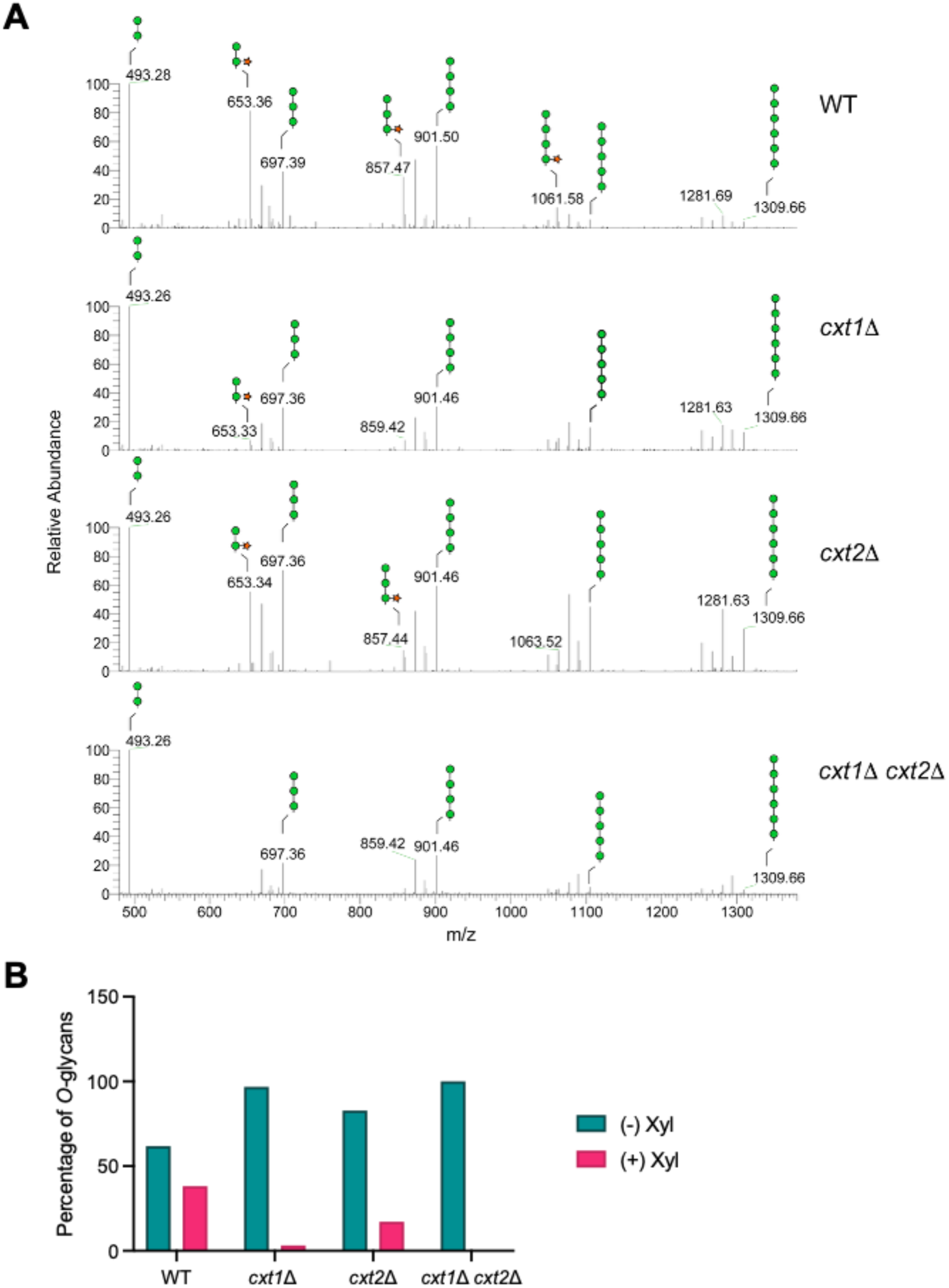
Cxt1 and Cxt2 mediate *O*-glycosylation. (A) Mass spectrometry spectra of *O*-linked glycans isolated from the indicated strains, showing the m/z region of the major *O*-linked glycans. Glycan structures are shown above the corresponding peak; specific m/z values and quantitation are in Table S1. Green circle, mannose; orange star, xylose. (B) Relative abundance of *O*-linked glycans with and without xylose.

### Both β-1,2-xylosyltransferases are needed for capsule synthesis

We previously found that Cxt1 acts in the synthesis of capsule polysaccharides: mutant strains exhibited an approximately 30% reduction of xylose in GXM and near-complete loss of mannose-linked xylose in GXMGal (41). However, we did not examine overall capsule size. To do this, we grew wild-type, *cxt1*Δ, *cxt2*Δ, and *cxt1*Δ *cxt2*Δ strains in “host-like” conditions that induce capsule expansion (here defined as DMEM tissue culture medium, 37°C, 5% CO_2_), where capsule synthetic enzymes might be most active (6, 49). We then visualized capsules by negative staining and measured their thickness. In JEC21, loss of either Cxt1, Cxt2, or both similarly reduced capsule size (Fig. 5A, C), suggesting that they play redundant roles in capsule synthesis in this background. Capsule imaging by transmission electron microscopy is consistent with this observation (Fig. S7A). Furthermore, the capsule fibers of the mutants appear twisted and short, similar to cells deficient in UDP-xylose synthesis or transport (36, 50). In KN99, both *cxt2*Δ and *cxt1*Δ *cxt2*Δ also had significantly smaller capsules (Fig. 5B, D). However, *cxt1*Δ capsules resembled those of the wild type, suggesting that Cxt2 is the predominant β-1,2-xylosyltransferase in capsule synthesis in this background. To further examine the physiochemical properties of these capsule polysaccharides, we immunoblotted shed capsule material after agarose gel electrophoresis, using the anti-GXM antibody 18B7 (Fig. S7B). GXM from JEC strains migrated similarly in all samples although *cxt1*Δ and *cxt1*Δ *cxt2*Δ had slightly stronger antibody binding. In KN99, both single mutants resembled WT, while *cxt1*Δ *cxt2*Δ exhibited altered migration and reduced antibody binding.

**FIG. 5.**
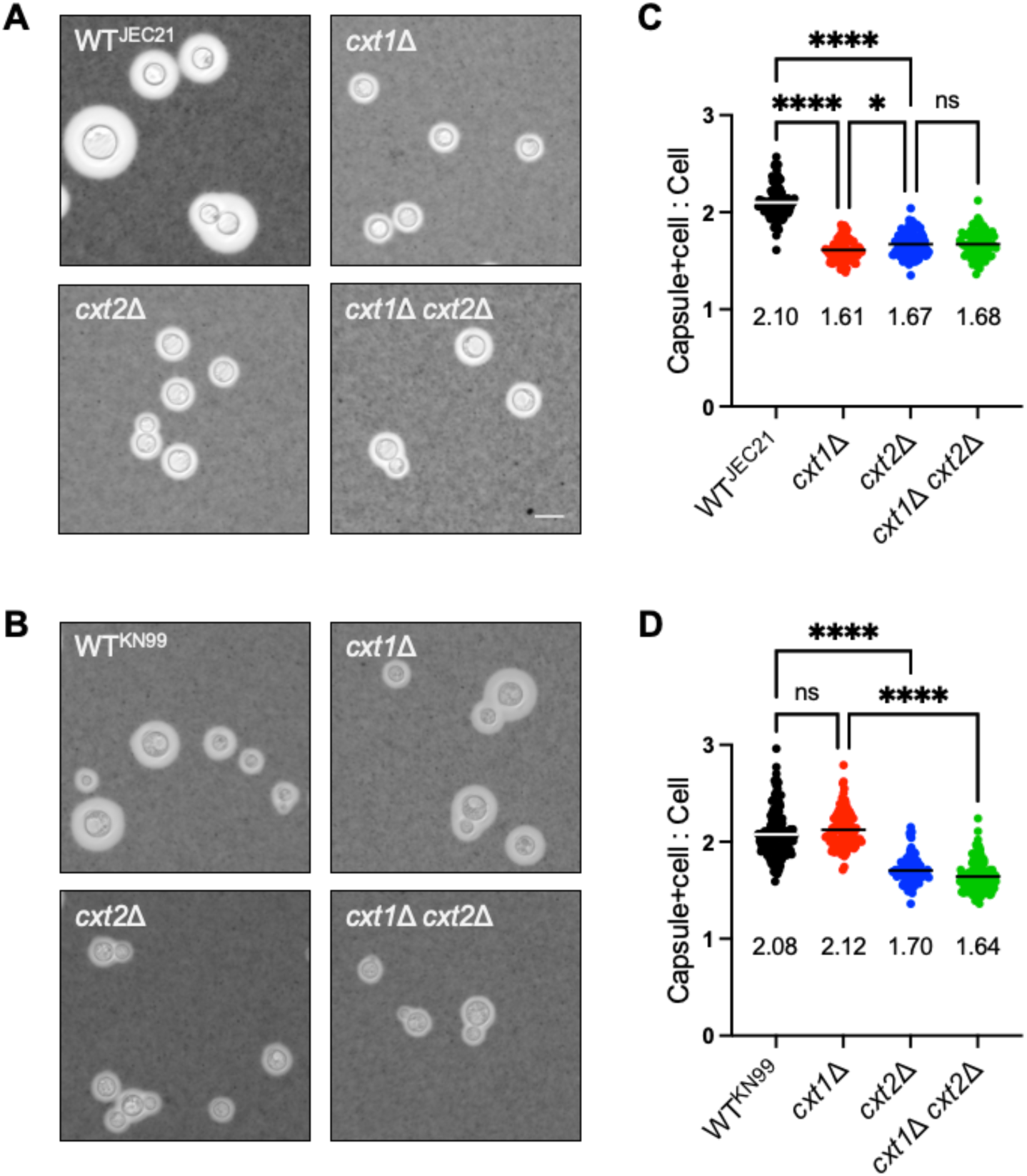
Cxt1 and Cxt2 influence capsule synthesis. (A-B) Representative images from India ink negative staining of JEC21 (A) and KN99 (B) strains grown in host-like conditions. All images are to the same scale; scale bar, 10 μm. (C-D) Quantitation of capsule sizes for the strains shown; plotted are values and mean for total radius (capsule + cell) relative to cell radius alone, for at least 80 cells. ****, p < 0.0001, *, p = 0.02 by ordinary one-way ANOVA with Tukey’s multiple comparisons test.

### β-1,2-xylosyltransferases impact virulence

Strains with smaller capsules are often defective in animal models of cryptococcal infection (9). Indeed, our results using an inhalational model of murine infection mirrored the observed capsule phenotypes. Strains in the overall less virulent JEC21 background showed lower lung burden for all three mutants (Fig. 6A), each of which has reduced capsule, with greater reduction in the strains that also display stress sensitivity (*cxt2*Δ and the double mutant; Fig. 2 and S4). The corresponding KN99 strains showed more subtle reduction, although it also correlated with decreased capsule size (Fig. 6B). Together, these results suggest that Cxt2 plays a role in virulence (see Discussion)

**FIG. 6.**
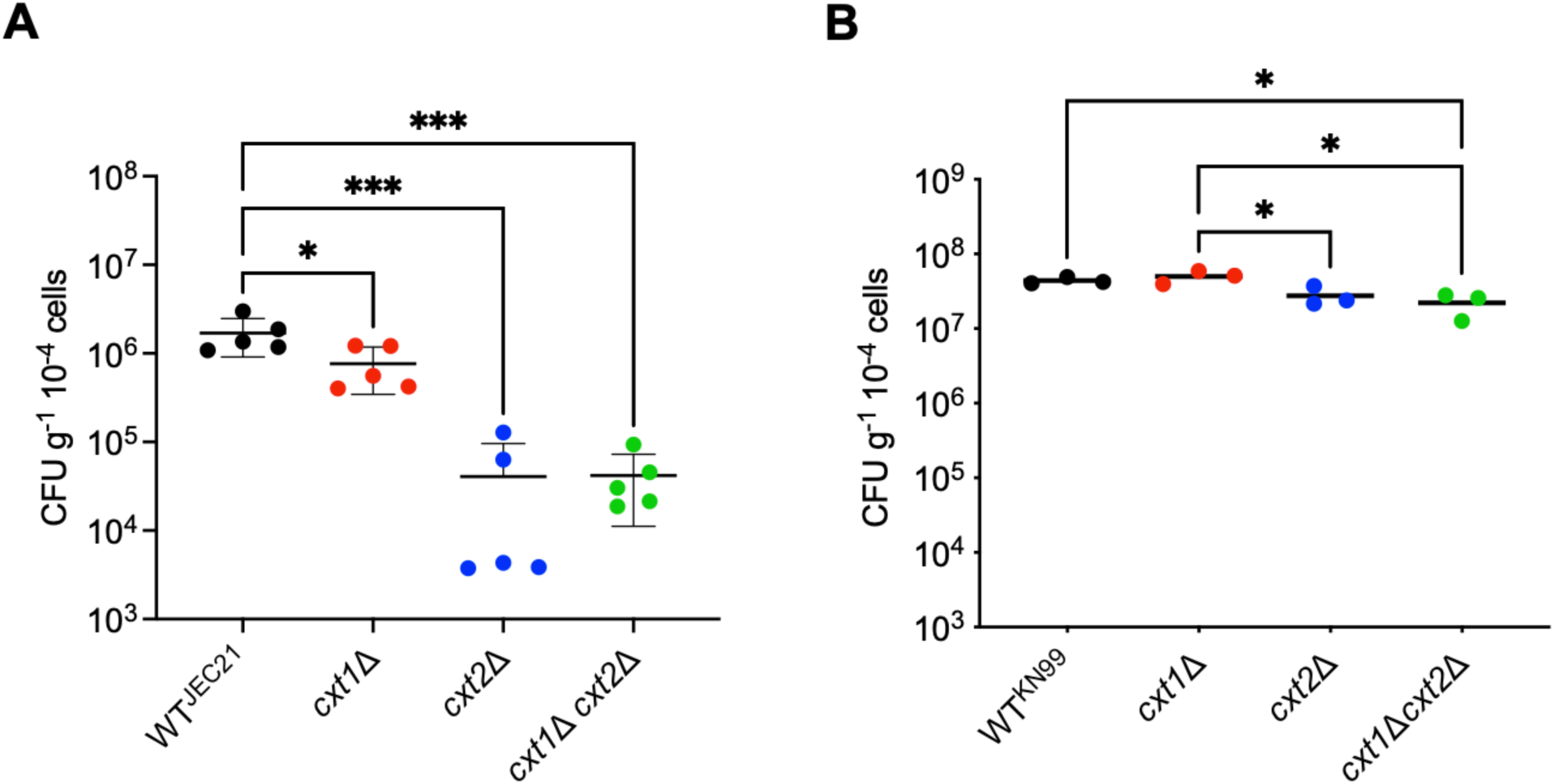
Impact of β-1,2-xylosyltransferases on virulence. Shown is lung burden of C57BL/6 mice sacrificed 12 days after intranasal infection with (A) 2.0×104 cells in the JEC21 background or (B) 2.5×104 cells in the KN99 background. Plots show the mean ± SD; each symbol represents one mouse. *, p < 0.05 and ***, p ≤ 0.0001 by ordinary one-way ANOVA with Tukey’s multiple comparisons test. Any comparisons not shown were not significant.

### Cxt1 and Cxt2 co-localize in the Golgi apparatus

Our results show that Cxt1 and Cxt2 participate in multiple glycosylation processes that occur in the secretory pathway, including GIPC synthesis, protein glycosylation, and capsule polysaccharide synthesis (24, 51–54). Several of these are initiated in the endoplasmic reticulum and proceed in the Golgi apparatus (24); we previously found that UDP-xylose transporters, which provide the precursor for xylose addition, occur in both compartments (36). To investigate the subcellular localization of Cxt1 and Cxt2, we epitope-tagged each protein. By immunofluorescence microscopy, Cxt1-Myc, expressed at the native locus under its endogenous promoter, colocalized with the GDP-mannose transporter Gmt1 (36, 55), indicating Golgi localization (Fig. 7A). We also compared Cxt1 localization to that of Cxt2-HA, driven by the actin promoter. Both proteins localized in large puncta distributed throughout the cytosol; in most of these puncta the tagged proteins colocalized (Fig. 7B). We also occasionally saw Cxt2 signal as a peri-nuclear ring, suggestive of ER localization (Fig. 7B, white arrowhead), although this may reflect the over-expression required for detection.

**FIG. 7.**
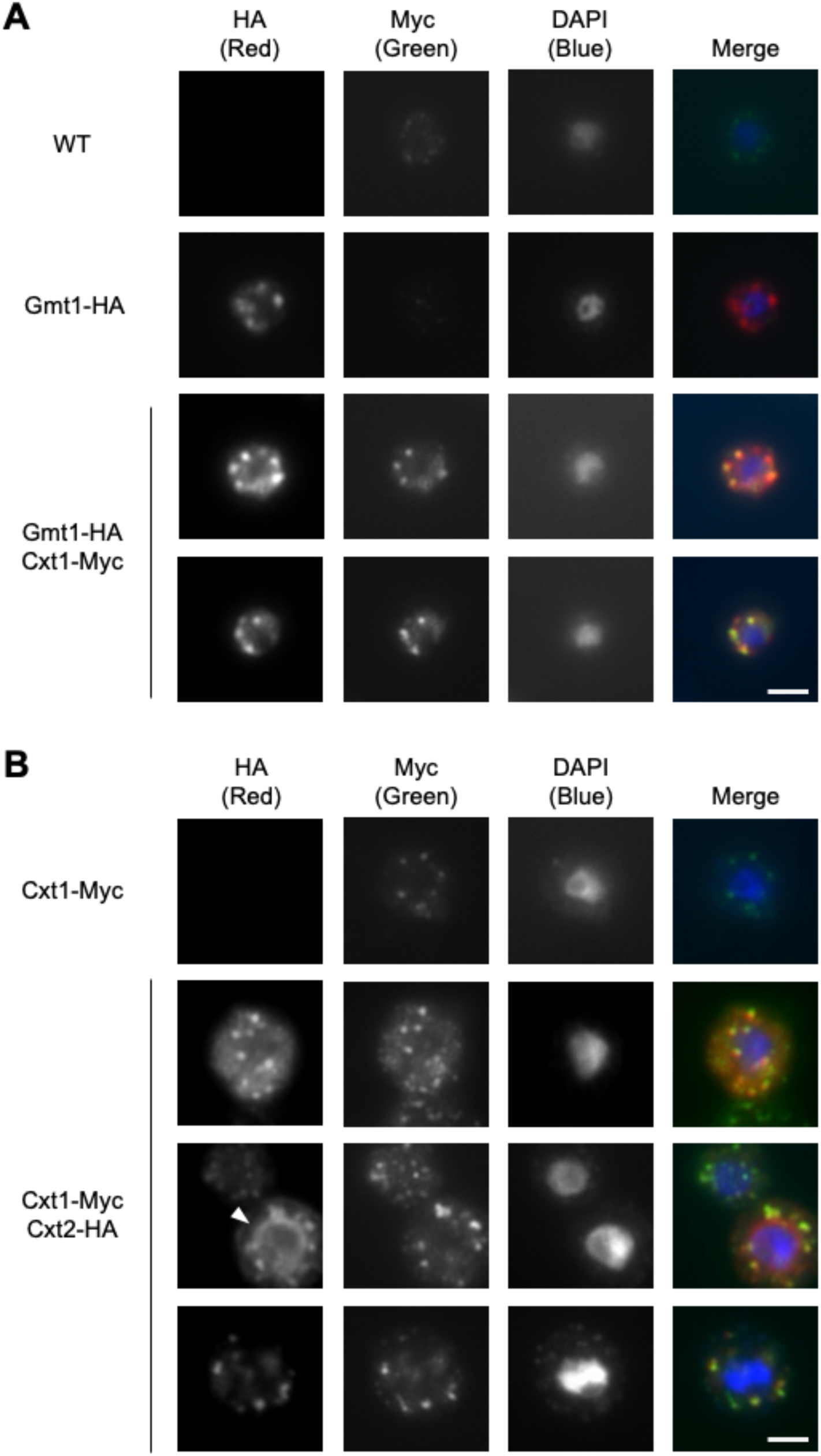
Localization of Gmt1, Cxt1, and Cxt2. Shown are representative immunofluorescence images of the indicated tagged strains in JEC21. Nuclei are stained with DAPI and the white arrowhead indicates peri-nuclear signal. All images are to the same scale; scale bar, 2 μm.

### Cxt1 and Cxt2 co-localize in the Golgi apparatus

Our results show that Cxt1 and Cxt2 participate in multiple glycosylation processes that occur in the secretory pathway, including GIPC synthesis, protein glycosylation, and capsule polysaccharide synthesis (24, 51–54). Several of these are initiated in the endoplasmic reticulum and proceed in the Golgi apparatus (24); we previously found that UDP-xylose transporters, which provide the precursor for xylose addition, occur in both compartments (36). To investigate the subcellular localization of Cxt1 and Cxt2, we epitope-tagged each protein. By immunofluorescence microscopy, Cxt1-Myc, expressed at the native locus under its endogenous promoter, colocalized with the GDP-mannose transporter Gmt1 (36, 55), indicating Golgi localization (Fig. 7A). We also compared Cxt1 localization to that of Cxt2-HA, driven by the actin promoter. Both proteins localized in large puncta distributed throughout the cytosol; in most of these puncta the tagged proteins colocalized (Fig. 7B). We also occasionally saw Cxt2 signal as a peri-nuclear ring, suggestive of ER localization (Fig. 7B, white arrowhead), although this may reflect the over-expression required for detection.

### Cxt1 and Cxt2 do not interact

There is precedent in yeast for protein-protein interactions between glycosyltransferases, which may contribute to the efficiency of glycoconjugate synthesis (56, 57). Given that Cxt1 and Cxt2 colocalize, we wondered if they might physically interact. To test this, we performed coimmunoprecipitation studies with a strain expressing both Cxt1-HA and Cxt2-FLAG, each under its native promoter. However, we found no evidence of interaction between these proteins, whether cells were grown in YPD or host-like conditions (Fig. S8).

### Cxt1 and Cxt2 are differentially expressed

Differential expression might also explain the distinct roles of Cxt1 and Cxt2. We previously published RNA-seq expression data for cells grown in host-like conditions over time (58). Analysis of this dataset showed a modest increase over time in expression of *CXT2*, with little change in that of *CXT1* (Fig. S9A). When we directly examined protein in the same conditions, we found that Cxt2-FLAG levels remained relatively steady while Cxt1-HA decreased over time (Fig. 8A, Fig. S9B). This results in a relative increase in the ratio of the two signals over time (Fig. 8B) (see Discussion).

**FIG. 8.**
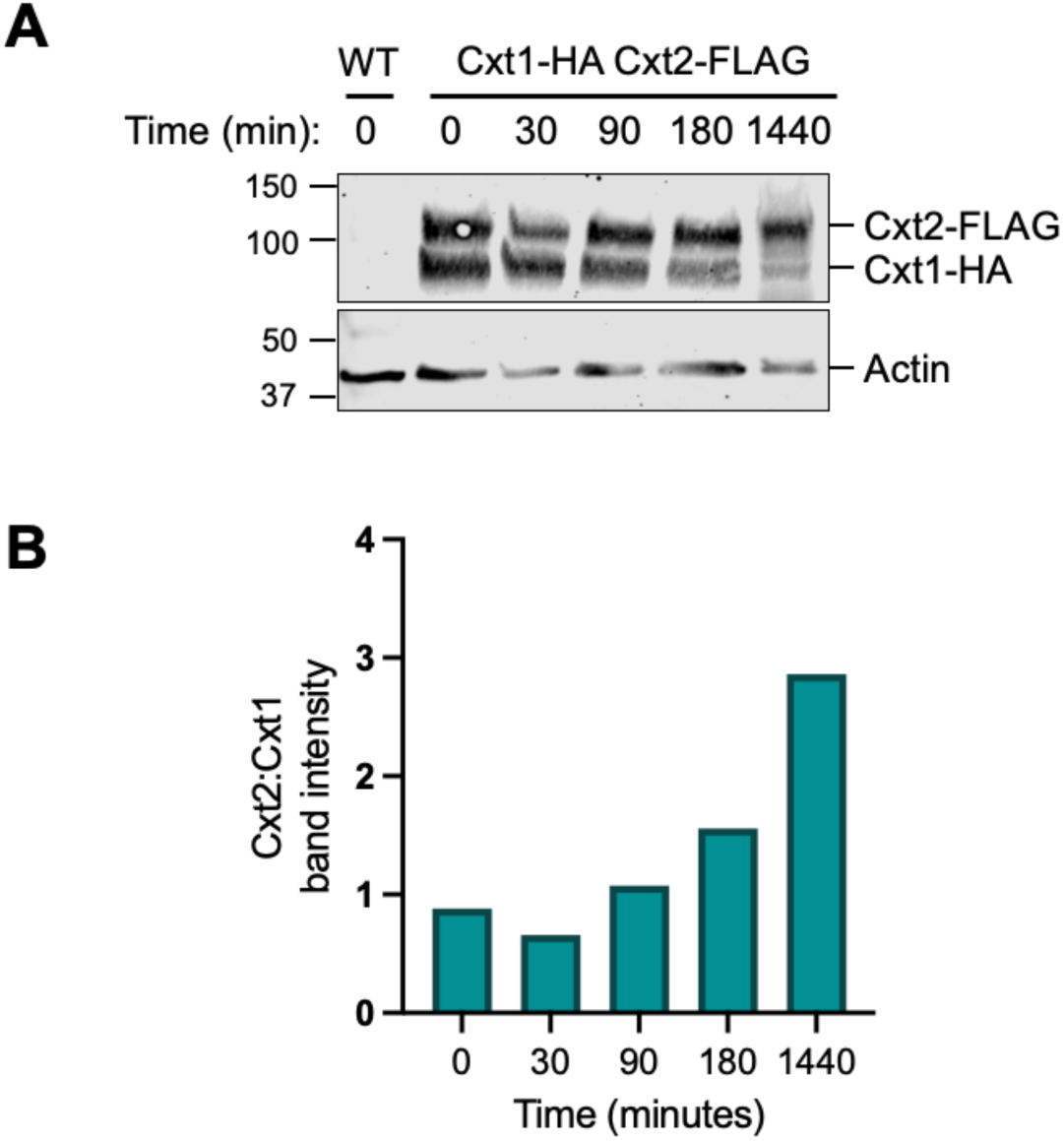
Gene and protein expression. (A) Immunoblot of protein lysates from wild-type and Cxt1-HA Cxt2-FLAG strains using anti-HA (1:5,000), anti-FLAG (1:500), and anti-actin (1:1,000) antibodies. 30 μg protein loaded per lane. MW markers indicated at left in kDa. Expected sizes: Cxt2-FLAG, 95 kDa; Cxt1-HA, 83 kDa; actin, 42 kDa. (B) Ratio of band intensity normalized to actin for Cxt2:Cxt1. The immunoblot and quantitation shown are representative of two experiments, each with two independent strains.

## DISCUSSION

The unique glycans of *Cryptococcus* play important roles in its biology and virulence. In previous studies, we identified Cxt1 as a β-1,2-xylosyltransferase that acts in the synthesis of GIPCs and capsule polysaccharides (33, 34, 41). Here, we have identified a second β-1,2-xylosyltransferase, Cxt2 (Fig. 1), and probed the biological roles of both enzymes.

Our most striking finding was that Cxt1 and Cxt2 together mediate all detectable xylose addition to *O*-glycans (Fig. 4), with Cxt1 playing the larger role. *O*-glycosylation is a highly conserved post-translational modification (59) with roles in protein folding, function, quality control, localization, and secretion (60–64). This process starts in the ER, where mannose is transferred from dolichol-phosphate-mannose to serine or threonine residues on polypeptides (65). In the Golgi, *O*-glycans are further extended by mannosyltransferases, forming linear structures of 1-6 α-mannose (up to 6 residues in our studies). In *Cryptococcus*, xylose is added to the first mannose residue at the reducing end of some of these *O*-mannosylated glycans (35). Cxt1 and Cxt2, which both localize in the Golgi (Fig. 7), are well-positioned to participate in this process.

Inhibition of *O*-mannosylation activates the cell wall integrity pathway, unfolded protein response, and ER-associated protein degradation (61). In *Cryptococcus*, mutants defective in *O*-linked extension show reduced cell wall integrity (CWI) signaling, stress resistance, virulence, and induction of the host immune response (35, 47). The phenotypic defects in *cxt1*Δ and *cxt2*Δ are more subtle, commensurate with the loss of only one residue.

Multiple studies report β-1,2-linked xylose in cryptococcal *N*-glycans (34, 38, 66), but neither Cxt1 nor Cxt2 seems to be directly involved in this process (Fig. 4) (34). We did find subtle differences at the population level (Fig. S6), but these changes may be indirect. For example, it may be that the absence of both Cxt1 and Cxt2 increases availability of UDP-xylose, the donor for xylose addition to *N*-glycans, explaining the higher fraction of *N*-glycans with this modification.

The polysaccharide capsule is a key feature in cryptococcal virulence. Cxt1 and Cxt2 are the first glycosyltransferases implicated in capsule synthesis to be localized, and are both found in the Golgi (Fig. 7). This supports our model of capsule synthesis occurring in that compartment (23). Cxt2 impacts capsule thickness in both strain backgrounds analyzed, while Cxt1 has less of a role in KN99 strains (Fig. 5). This corresponds with virulence in a mouse model of infection, where mutants lacking *CXT2* demonstrate reduced virulence relative to wild type (Fig. 6). Overall, our findings suggest that Cxt2 is more important for virulence than Cxt1.

Some mutant phenotypes, such as stress resistance (Fig. 2, S4), differ by serotype. JEC21 strains unsurprisingly exhibit more severe defects, as *C. deneoformans* is generally less virulent and thermotolerant relative to *C. neoformans* (19, 20). Other reports have also noted variation in phenotypes depending on strain background (67, 68). It is interesting that although the Cxt proteins have similar biochemical activity in both serotypes, the strain background ultimately dictates their functional impact on cell growth and virulence.

Some phenotypes we observed rely predominantly on one Cxt protein, as with the dependence of GIPC modification on Cxt1 (Fig. S5). For others, deletion of either gene yields a similar phenotype, as for capsule size in JEC21 (Fig. 5A, C). In still others, loss of both proteins is required to manifest the maximal phenotypic defect, as with *O*-glycosylation (Fig. 4). Together, these results suggest that the two proteins play distinct yet overlapping roles. This raised the questions of their relative expression and potential interaction, which we also pursued.

Analysis of gene expression over time, using our previously published data (58), suggested that while *CXT1* is the more highly expressed of the two genes (Fig. S9A), *CXT2* expression increases over time in host-like conditions, further supporting its role in virulence. RNA-seq of JEC21 grown at 30°C vs. 37°C by Wallace *et al* revealed similar patterns of expression (69). Interestingly, protein expression does not keep pace with gene expression for either one, with Cxt2 holding steady over time while Cxt1 decreases (Fig. 8A). This could be due to mRNA stability, processing, or efficiency of translation. (Protein amounts cannot be directly compared, because different antibodies were used to detect them.) Despite their colocalization, we did not detect any direct interactions between Cxt1 and Cxt2, although they may interact indirectly or transiently (Fig. S8).

Overall, we found that Cxt1 and Cxt2 impact xylose addition to multiple major cryptococcal glycoconjugates, including GXM, GXMGal, GIPCs, and protein *O*-glycans, but are not directly involved in synthesis of protein *N*-glycans. This suggests that *C. neoformans* encodes at least one other β-1,2-xylosyltransferase. Although we did not detect such an enzyme in our *in vitro* assays (Fig. 1), it may be lowly expressed in the rich growth conditions used for these assays or require a different acceptor than the α-1,3-mannobiose provided.

Xylose modification of proteins is known to impact the human immune response. In the model plant *Arabidopsis*, decreased exposure of xylose on *N*-glycans leads to reduced immunogenicity in allergy patients (70). In *Cryptococcus*, mutants with a higher proportion of xylose-containing *O*-glycans have enhanced immunogenicity relative to wild-type strains, and vice versa (39). Conversely, loss of UDP-xylose synthesis (and thus xylose modification) leads to increased uptake by host macrophages (71) and enhanced complement activation (72). Strain-dependent variation in cryptococcal capsule composition, much of which involves altered xylose modification, also elicits different immune responses (73). Although uptake of *cxt1*Δ and *cxt2*Δ by host macrophages is unaltered relative to wild type (71), it will be interesting to investigate the impact of xylose modification on other aspects of the host immune response.

In this study, we examined the enzymatic activity and biological relevance of two β-1,2-xylosyltransferases, Cxt1 and the newly discovered Cxt2. These two enzymes play overlapping but distinct roles in cryptococcal biology, which interestingly differ between cryptococcal serotypes, highlighting the importance of utilizing multiple strain backgrounds. Our experiments resolve previously unknown components of cryptococcal glycan synthesis and identify the enzymes responsible for *O*-glycan xylose modification, moving us closer to a deep understanding of synthesis of the unique glycoconjugates of *Cryptococcus*.

## METHODS

### Strain construction and growth

Strains used in this study are listed in Table S2 and their construction is detailed in the Supplemental Text. All strains used in this study are mating type alpha. For both *C. neoformans* and *C. deneoformans* strain sets, deletion mutants and wild-type strains were isogenic products of mating. For all experiments, cryptococcal strains were grown on yeast extract-peptone-dextrose (YPD) plates for 2 days at 30°C. For liquid cultures, single colonies were inoculated into YPD and grown overnight at 30°C, with shaking. For capsule induction in host-like conditions, overnight YPD cultures were washed in PBS and grown in high-glucose DMEM (Sigma, D6429) at 10^6^ cells/mL for 24 h at 37°C and 5% CO_2_, without shaking.

### Crude membrane preparation and xylosyltransferase activity assays

Crude membrane preparations and xylosyltransferase activity assays were performed as we previously described (40). Overnight YPD cultures were resuspended in 100 mM Tris pH 8.0, 0.1 mM EDTA and disrupted by bead-beating. Unbroken cells and debris were removed by centrifugation (1,200 g; 20 min) and the resulting supernatant fractions were subjected to further centrifugation (60,000 g; 45 min). The 60,000 g pellets were thoroughly resuspended in the same buffer, the membranes resedimented in the same conditions, and the supernatants discarded. The washed membrane pellets were solubilized with 1% Triton X-100 on ice for 30 min with brief vortexing every 5 min, then centrifuged again (60,000 g; 30 m) and the supernatant fractions recovered as solubilized crude membranes.

To assay xylosyltransferase activity, solubilized crude membranes were incubated with 57 nmol UDP-[^14^C]xylose and 8.5 mM α-1,3-mannobiose for 4 h at 20°C. To remove excess UDP-[^14^C]xylose and terminate the reaction, the assay mixtures were applied to AG2X-50 resin and eluted with deionized water. The eluates were centrifuged (13,000 g; 5 min) and the supernatants, containing soluble reaction products, were collected. For Jack Bean α-mannosidase digestion, the products were treated for 18 h at 37°C as in reference (40). The ^14^C-labeled products were detected by thin-layer chromatography (TLC).

For TLC, reaction products were dried under nitrogen at 50°C, resuspended in 15 μL 40% *n*-propyl alcohol, and applied to 20 x 20 cm Silica Gel-60 TLC plates. Plates were developed in 5:4:2 *n*-propyl alcohol:acetone:water for 2 h, allowed to dry, and then developed again for 2 h. After drying, standards were visualized by spraying with 0.2% orcinol in 75:15:10 ethanol:sulfuric acid:water and heating at 70°C for 5-10 m. Sample lanes were sprayed with Enhance surface autoradiography spray and then visualized by autoradiography.

### Phenotyping

Overnight YPD-grown cultures were adjusted to 10^7^ cells/mL in PBS. 3 μL aliquots of 10-fold serial dilutions were spotted on YPD agar without or with 1.2 M NaCl, 0.01% SDS, 2.2 mM calcofluor white, or 10% DMSO (calcofluor white vehicle control). Plates were incubated at 30°C or 37°C for 72 h.

### Glycoprotein preparation, oligosaccharide release, and mass spectrometry

Cell lysates were prepared as we previously described (48), with all steps performed at 4°C with ice-cold buffer. Dense YPD cultures were sedimented and resuspended in 100 mM pH 8.0 Tris, 0.1 mM EDTA, and then disrupted by bead-beating. Samples were centrifuged (2,000 g; 25 m) to remove unbroken cells and debris, and the crude lysate (supernatant) was solubilized with 1% CHAPS (final concentration) by rotating for 2 h at 4°C. Insoluble material was removed by centrifugation (75,000 g; 45 min) and the extracts were dialyzed against 2 L ice-cold 50 mM ammonium bicarbonate buffer using 6-8 kDa Fisher brand regenerated cellulose dialysis tubing for 48 h (with buffer changes every 12 h). Dialyzed samples were lyophilized, then washed with ice-cold 80% acetone to remove residual detergent and other contaminants.

*O*-linked oligosaccharides were released by reductive β-elimination as and permethylated as we previously described (74). *N*-linked oligosaccharides were released from lyophilized cell extract material by PNGase F digestion and permethylated as we previously described (75, 76). Samples were then analyzed by mass spectrometry as detailed in the Supplemental Text.

Mass spectra for these studies are available in GlycoPOST under accession number: GPST000698 (77). Glycomics data and metadata were obtained and are presented in accordance with MIRAGE standards and the Athens Guidelines (78, 79). Symbol and text nomenclature for representation of glycan structures is according to the Symbol Nomenclature for Glycans (SNFG) (80). The explicit identities of individual monosaccharide residues are assigned based on known fungal biosynthetic pathways. Data annotation and assignment of glycan accession identifiers was facilitated by the GRITS Toolbox (81), GlyTouCan (82), and GlyGen (83, 84).

### Capsule analysis

Cell cultures were induced as described above, then washed, resuspended in PBS, and mixed with India ink (Higgins) in PBS (2:1 vol/vol) to visualize capsule by negative staining. Randomly chosen fields of cells were imaged on a ZEISS Axio Imager M2 fluorescence microscope. The areas of each cell body with and without capsule were measured by ImageJ; their radii were calculated for at least 80 cells per sample and plotted as a ratio.

### Virulence studies

All animal protocols were approved by the Washington University Institutional Animal Care and Use Committee, with care taken to minimize animal handling and discomfort. Groups of three or five 6-8-week-old female C57BL/6 mice from Jackson Laboratory were anesthetized by subcutaneous injection of 120 μL of 10mg/mL ketamine and 2 mg/mL xylazine in sterile PBS and intranasally infected with either 2.0 x 10^4^ cells (JEC21 background) or 2.5 x 10^4^ cells (KN99 background) in 50 μL PBS. Mice were monitored with care and humanely sacrificed 12 days post-infection; lungs and brains were harvested, homogenized, and plated on YPD agar to assess fungal organ burden. Colony-forming units (CFU) were counted, normalized to organ weight, and analyzed by ordinary one-way ANOVA with Tukey’s multiple comparisons test.

### Immunofluorescence

Immunofluorescence was performed as in reference (48). Briefly, cells were cultured overnight in minimal medium (0.17% w/v YNB without amino acids and (NH_4_)_2_SO_4_, 0.5% w/v (NH_4_)_2_SO_4_, 2% w/v glucose) at 30°C, fixed with 3.7% formaldehyde, and subjected to partial cell wall digestion before 1% Triton X-100 permeabilization, staining and imaging as detailed in the Supplemental Text.

### Protein extraction and immunoblotting

YPD-grown cells were transferred to high-glucose DMEM at 37°C and 5% CO_2_ and grown for 0, 30, 90, 180, or 1440 min before protein extraction. Cells were sedimented, washed in cold MilliQ water, and resuspended in ice-cold immunoprecipitation lysis buffer (100 mM Tris pH 7.6, 150 mM NaCl, 1 mM EDTA, 1% NP-40, and protease inhibitor cocktail (Sigma P8215)). Cells were next disrupted by bead-beating (8 rounds of 1 min bead beating followed by 1 min on ice) and unbroken cells were removed by centrifugation (500 g; 20 min; 4°C). The protein concentration of the resulting supernatant (cleared cell lysates) was determined by BCA (Pierce BCA Protein Assay Kit, 23225).

Protein samples were mixed with 5x SDS sample buffer containing beta-mercaptoethanol and boiled for 5 min. Samples were resolved on 4-15% SDS-PAGE gels and transferred to nitrocellulose membranes, which were blocked with 10% milk in TBS for 1 h, washed with TBS Tween-20, and incubated with primary antibodies (1:5,000 anti-HA antibody (Abcam, ab9110), 1:500 anti-FLAG (Millipore Sigma, F3165), and 1:1,000 anti-actin antibody (ThermoFisher, MA1-744)) overnight at 4°C. Membranes were washed with TBS Tween-20, incubated with secondary antibody (1:10,000 680RD goat anti-mouse IgG (LI-COR IRDye, 926-68070) or 1:10,000 800CW goat anti-rabbit IgG (LI-COR IRDye, 926-32211)) for 1 h at room temperature, washed with TBS, and visualized on a LI-COR 92120 Odyssey Infrared imaging system.

### Statistical methods

Statistical analysis was performed using Prism (GraphPad) for ordinary one-way ANOVA with Tukey’s multiple comparisons test.

### Additional methods

Detailed methods for RNA interference, Protein sequence alignment and structure analysis, Glycolipid preparation and analysis, and Co-immunoprecipitation are in the Supplemental Text.

## ACKNOWLEDGEMENTS

We appreciate the generosity of Guilhem Janbon in supporting the naming of CKF44_01283 as *CXT2* to align with the demonstrated function of its protein product, although he had earlier called the corresponding H99 gene *CAP5* based on homology. We are grateful to the late Steven B. Levery for GIPC analysis. We also thank Morgann Reilly and Zhuo Wang for protein preparation, Matthew Williams for assistance with animal studies, Hong Liu and Cara Griffith for cloning, Dr. Wandy Beatty (Washington University School of Medicine) for TEM, and Dr. Thomas R. Kozel (University of Nevada School of Medicine) and Dr. Arturo Casadevall (Johns Hopkins University) for anti-GXM monoclonal antibodies. We appreciate helpful discussion and comments on the manuscript from Sydney L. Briner, Zanetta Chang, Cecilia Gutierrez-Perez, Thomas Hurtaux, and Dong-gyu Kim. We also acknowledge assistance on this project from former Washington University undergraduate students Matt Vogt, Alyssa Marulli, Sunday Oladipopo, and Katie Lewandowski.

These studies were supported by NIH R01 awards AI049173, AI135012, and AI192892 to TLD. DBK was partly supported by NIH T32 GM139774. Deconvolution of mass spectrometry data was supported by the Mass Spectrometry Core, Translational Metabolomics Shared Resource (RRID: SCR_027908).

