## Supplemental Material for "Two closely related β-1,2-xylosyltransferases differentially impact cryptococcal glycan synthesis"

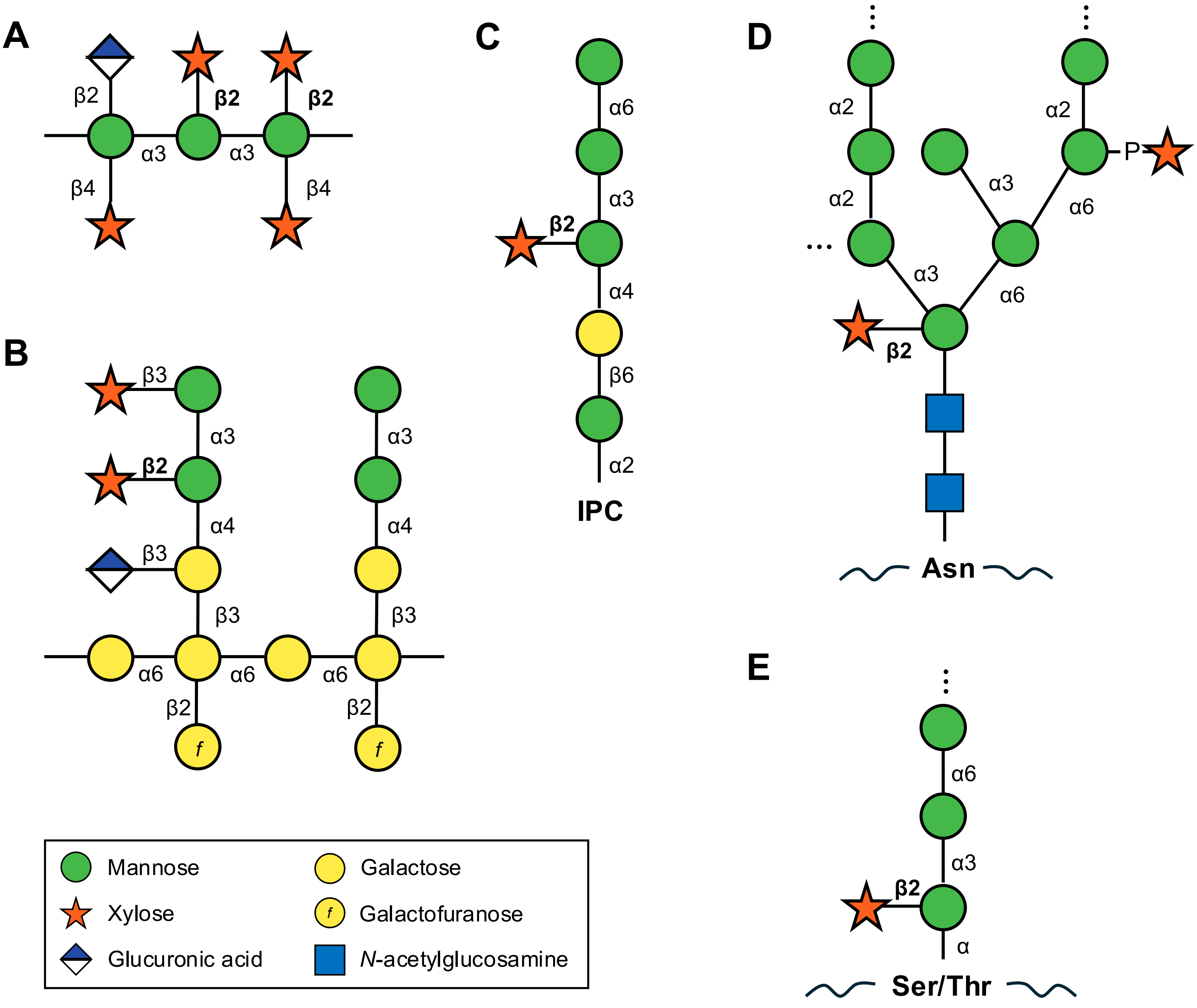


**FIG. S1.** Examples of xylose-containing glycoconjugates in *Cryptococcus*: (A) the repeating subunit of capsule GXM, (B) the repeating subunit of capsule GXMGal, (C) glycosylinositol phosphorylceramides, (D) *N*-glycans, and (E) *O*-glycans. Linkages of xylose β-1,2 to the reducing mannose of α-1,3-mannobiose are in bold. IPC, inositol phosphorylceramide; Asn, asparagine; P, phosphate; Ser, serine; Thr, threonine.

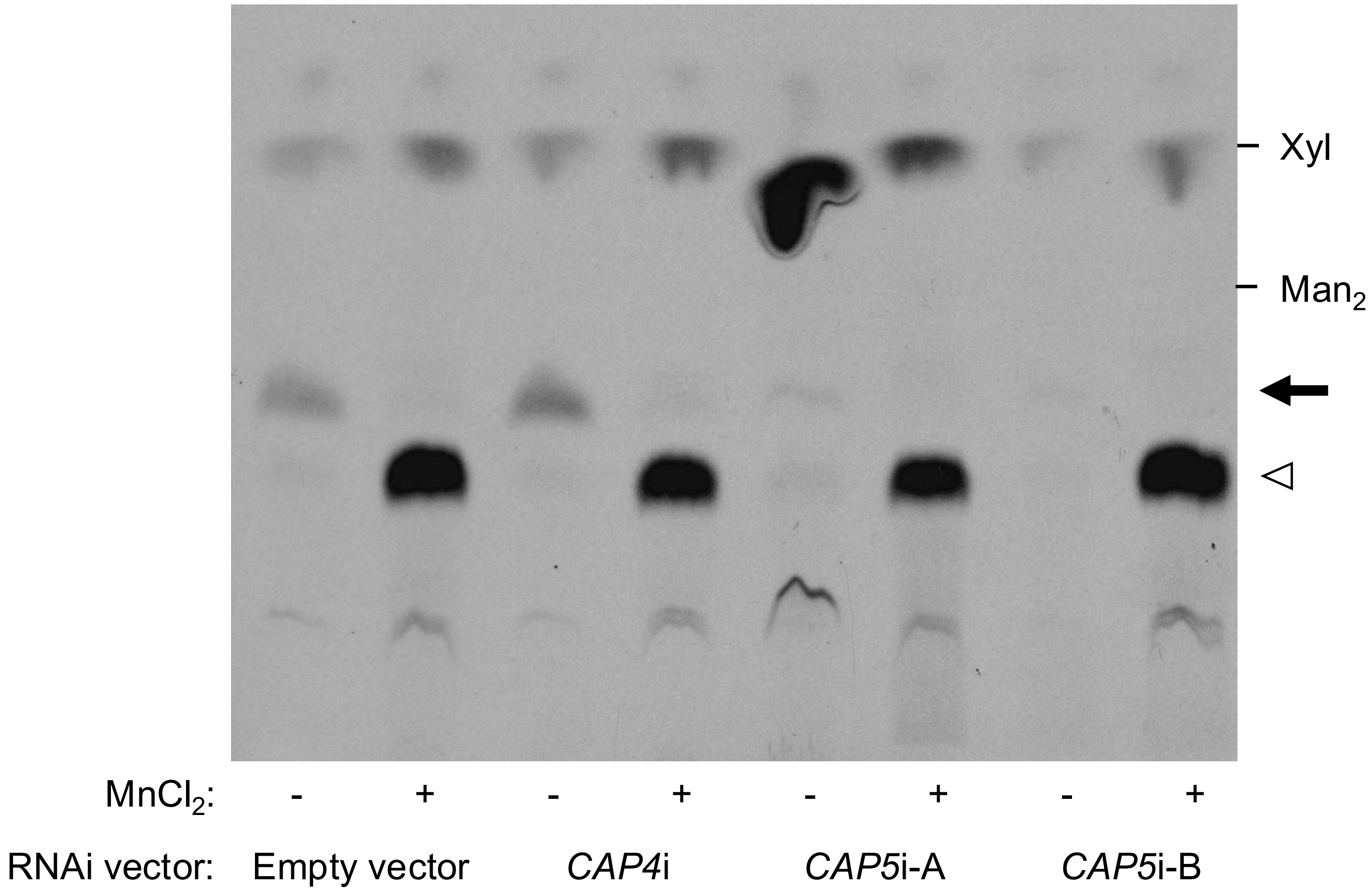


**FIG. S2.** RNAi reveals a second β-1,2-xylosyltransferase. Autoradiographs of β-1,2-xylosyltransferase assay products resolved by TLC. Assays were performed using washed membranes from JEC21 cells transformed with empty vector or with vector including sequences designed to silence *CAP4* (*CAP4i*) or *CAP5* (*CAP5i-A* and *CAP5i–B*, which target independent sequences). Parallel assays were performed under standard assay conditions (no $MnCl_2$) and in the presence of $MnCl_2$. The latter stimulates formation of an unrelated product (Xyl-phosphate-$Man_2$ (Supp. Text ref. 1); open arrowhead), which out-competes the Cxt1 activity for radiolabeled substrate; this serves as an internal control for enzymatically active membranes. Migration positions of non-radiolabeled standards are indicated at the right; the black arrow indicates the migration position of the Cxt1 product.

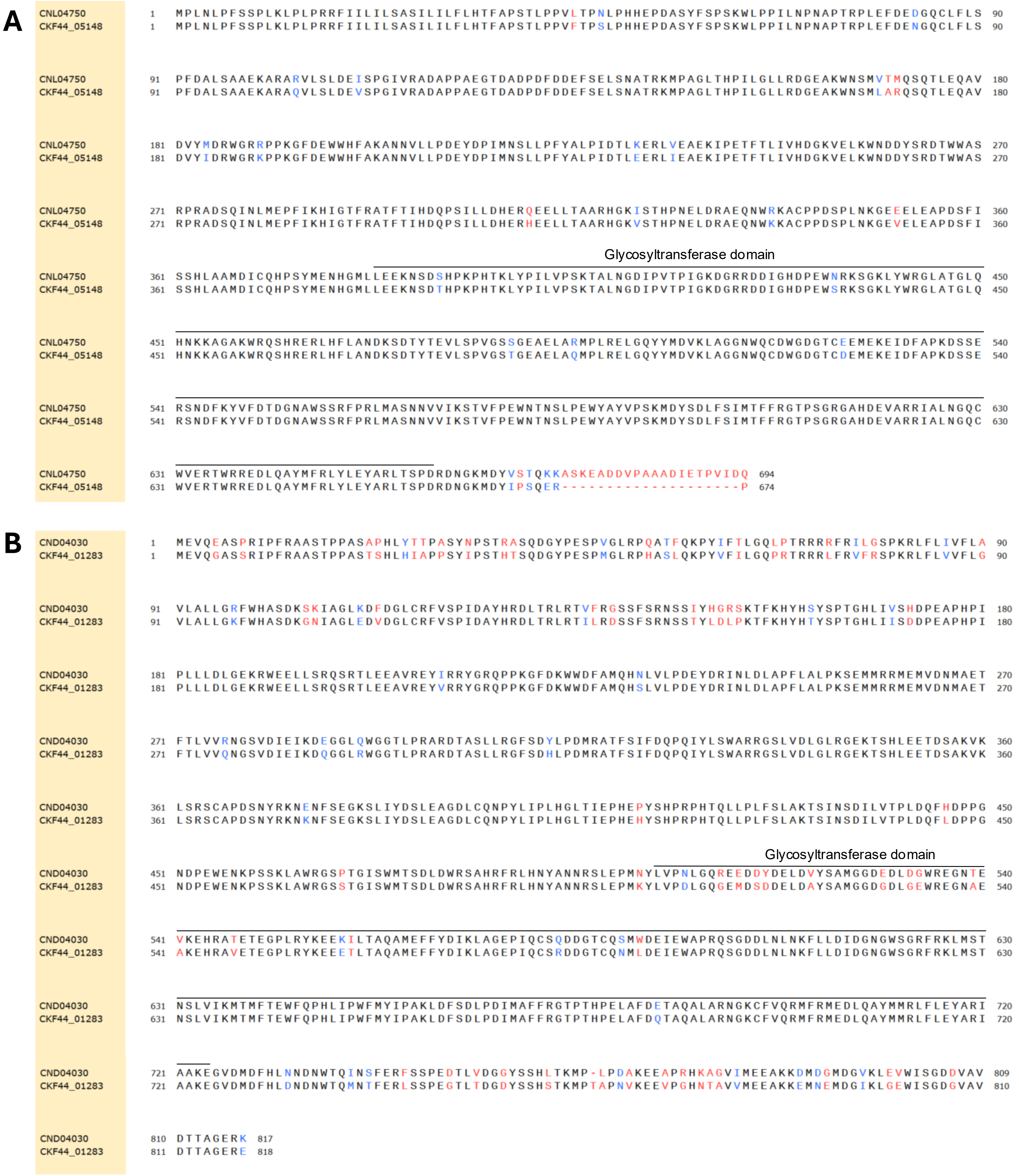


**FIG. S3.** Conservation of cryptococcal Cxt1 and Cxt2. (A and B) NCBI Protein BLAST sequence alignment of Cxt1 (A; JEC21 CNL04750 and KN99 CKF44_05148) and Cxt2 (B; JEC21 CND04030 and KN99 CKF44_01283). Black residues, identical; blue, similar; red, not similar. Glycosyltransferase domains are marked with InterProScan and SMART predictions according to FungiDB annotations (Supp. Text ref. 2) (continued on next page)

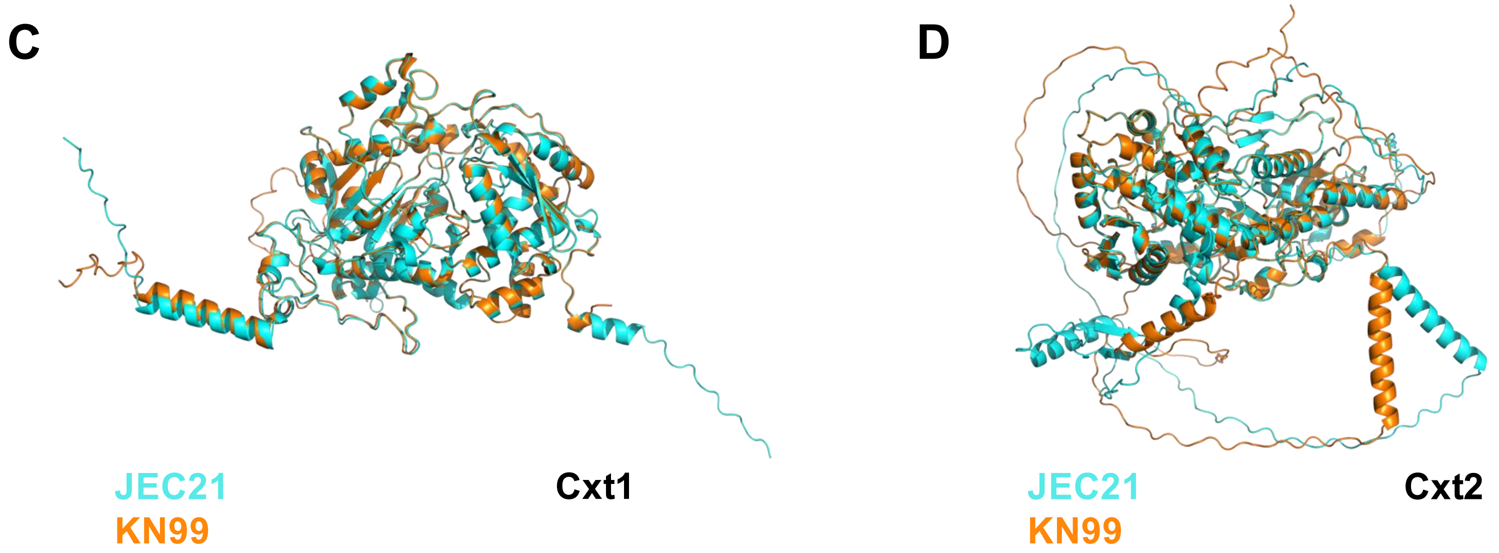


**FIG. S3.** (continued) (C and D) Structure alignment using AlphaFold predicted structures superimposed on Pymol. JEC21 (cyan) and KN99 (orange) sequences are shown for Cxt1 (C) and Cxt2 (D).

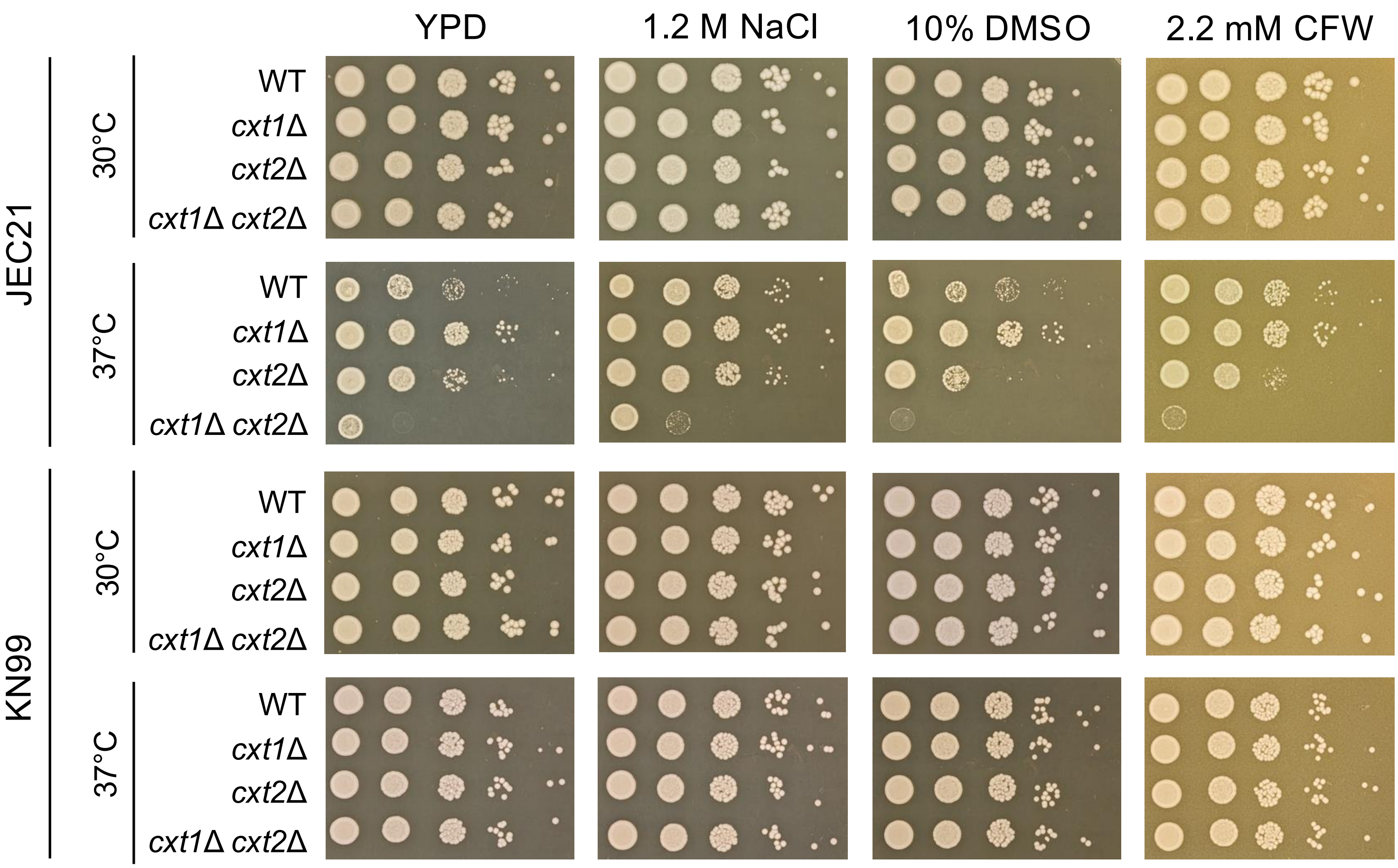


**FIG. S4.** Impact of β-1,2-xylosyltransferases on growth *in vitro*. The indicated strains were plated in 10-fold serial dilutions on YPD agar medium -/+ 1.2 M NaCl, 0.01% SDS, 10% v/v DMSO (vehicle control for calcofluor white), or 2.2 mM calcofluor white (CFW) and incubated at 30°C and 37°C for 72 h as shown. These plates were from the same experiment shown in Fig. 2, so the same YPD plates are included as controls here. The experiment shown is representative of three similar studies.

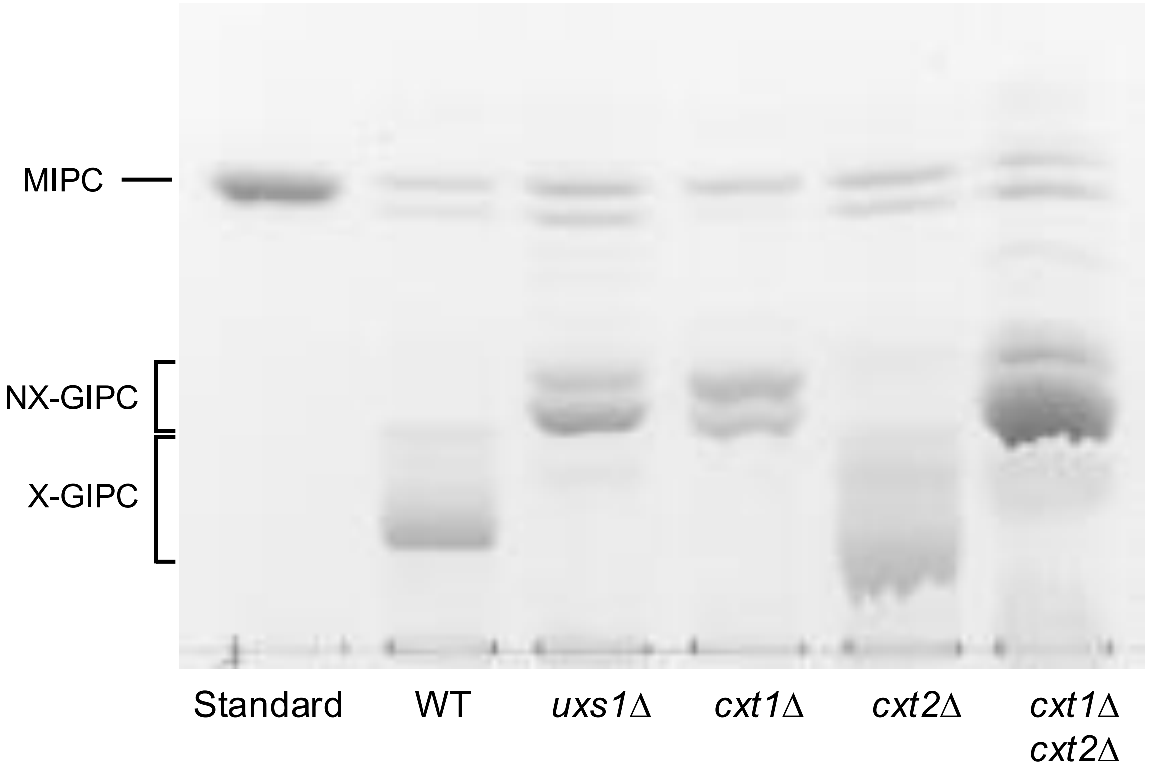


**FIG. S5.** Cxt1 but not Cxt2 impacts GIPC synthesis. GIPCs were isolated from washed cell membranes of the indicated *C. deneoformans* strains, resolved on HPTLC, and stained with orcinol as described in the Supplementary Methods. MIPC, mannosylinositol phosphorylceramide standard (Supp. Text ref. 3); NX-GIPC, non-xylosylated GIPC; X-GIPC, xylosylated GIPC.

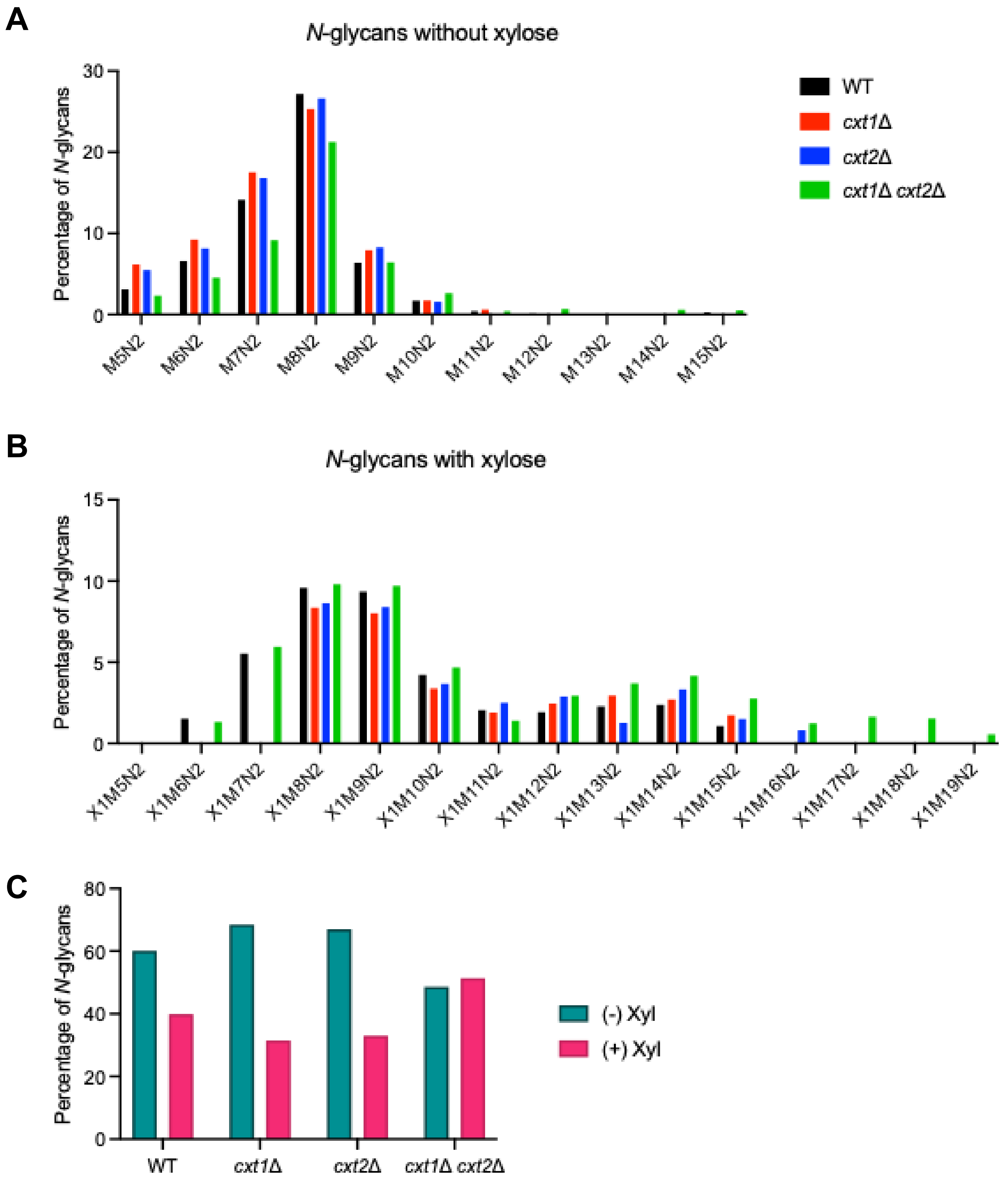


**FIG. S6.** Relative quantitation of *N*-glycan abundance for the indicated strains. Shown is the distribution of *N*-glycan structures without (A) and with (B) xylose, as a percentage of total *N*-glycans. Structures are indicated by the number of xylose (X), mannose (M), and *N*-acetylglucosamine (N) residues. (C) Percentage of *N*-glycans -/+ xylose.

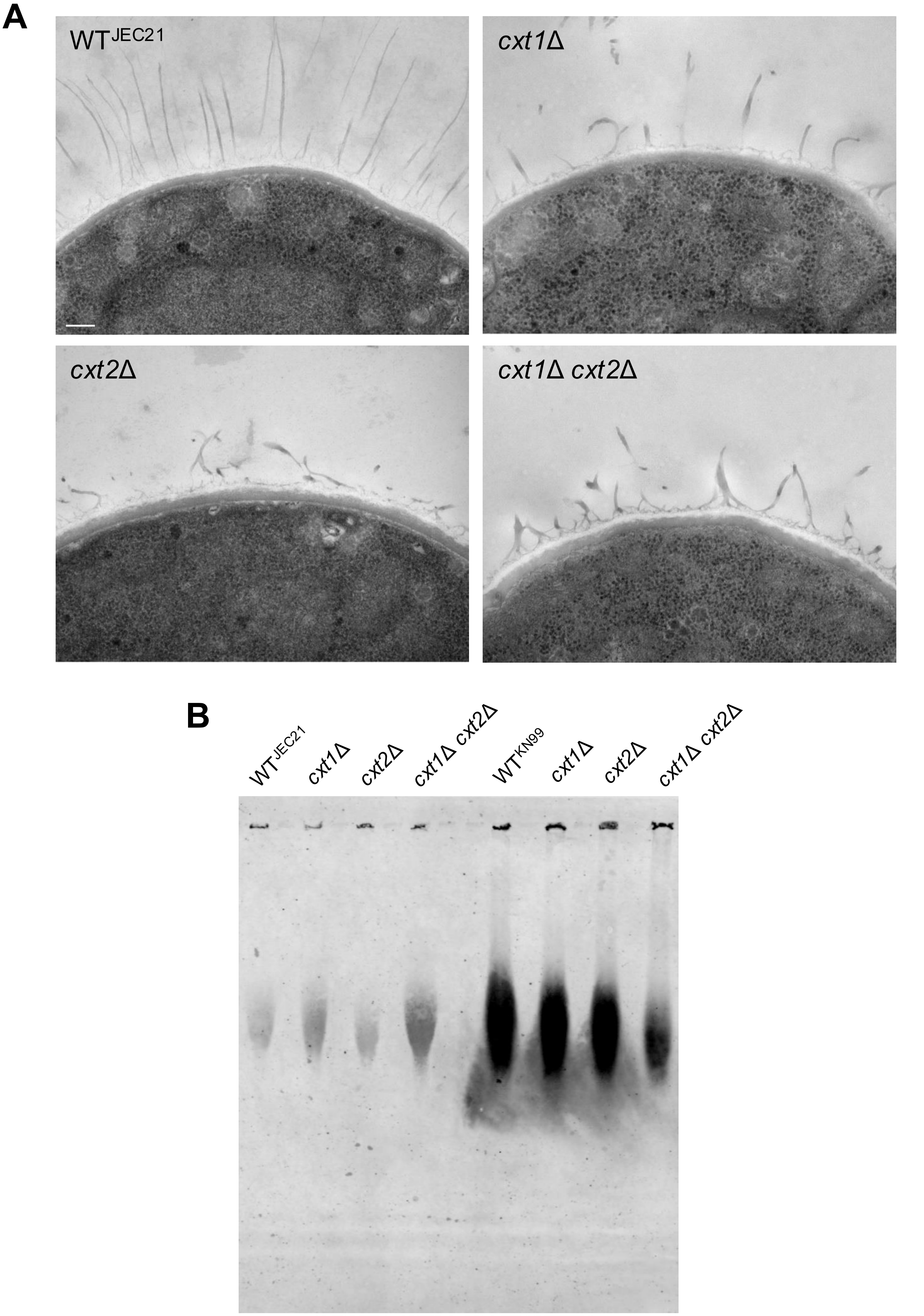


**FIG. S7.** Cxt1 and Cxt2 influence capsule synthesis. (A) Representative transmission electron micrographs of JEC21 strains grown in YPD. All images are to the same scale; scale bar, 200 nm. (B) GXM immunoblot of dialyzed conditioned media from the indicated strains, probed with anti-GXM antibody 18B7.

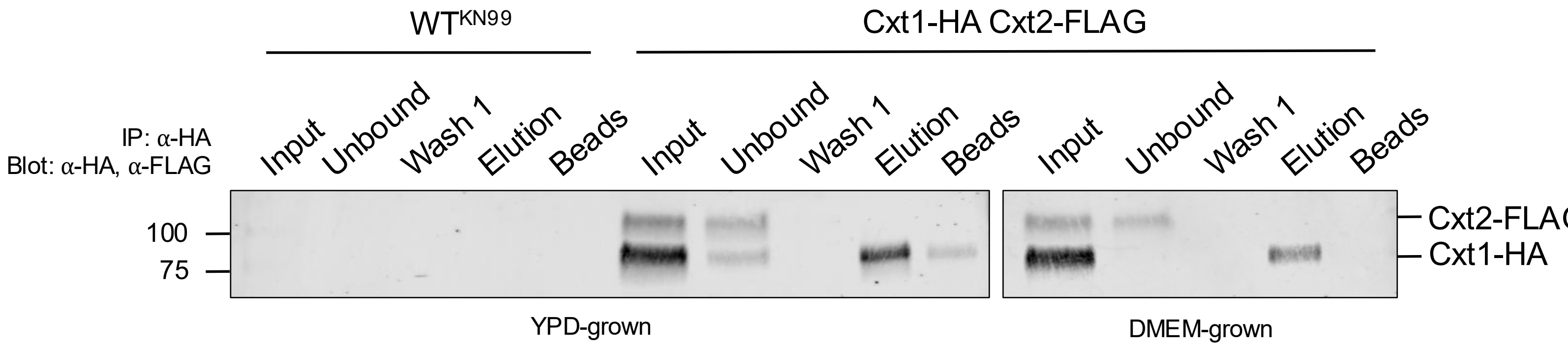


**FIG. S8.** Cxt2 does not co-immunoprecipitate with Cxt1. 500 µg of detergent extracts from WT or epitope-tagged strains grown in YPD at 30°C overnight or DMEM for 30 min at 37°C, 5% $CO_2$ (Input) were incubated with anti-HA beads as in the Supplementary Methods. Beads were separated from unbound material by centrifugation, washed, and heated in 2x sample buffer to elute bound proteins. Equal proportions of each fraction were resolved by 4-15% gradient SDS-PAGE electrophoresis and subjected to immunoblotting with a combination of anti-FLAG (1:2,000) and anti-HA (1:5,000) antibodies. Expected sizes: Cxt2-FLAG, 95 kDa; Cxt1-HA, 83 kDa. Results are representative of those obtained with two independent tagged strains.

A

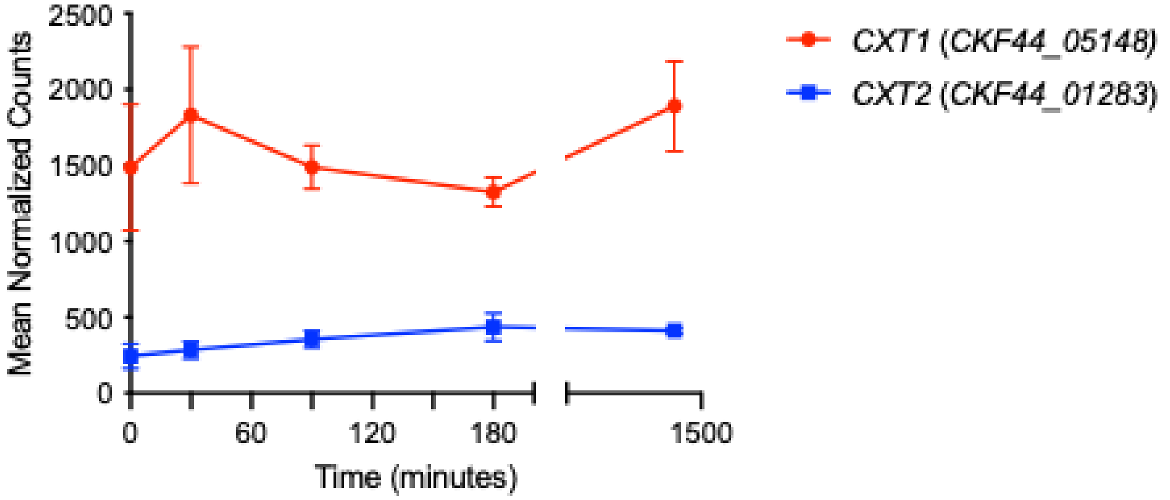


B

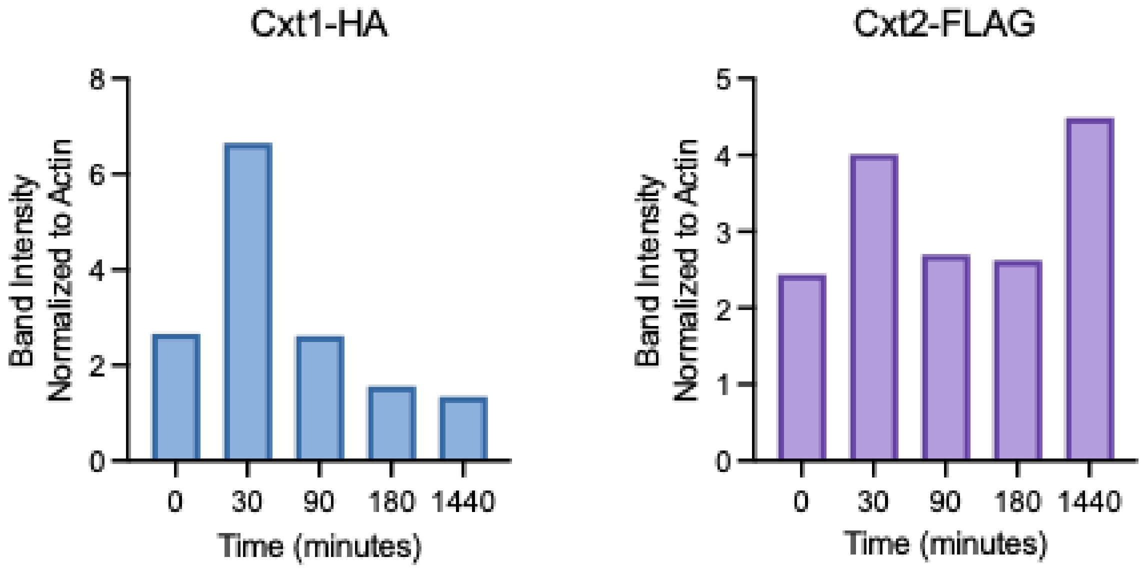


**FIG. S9.** Cxt1 and Cxt2 expression. (A) Expression of *CXT1* and *CXT2* in KN99 cells grown in DMEM, 37°C, 5% $CO_2$ over time, plotted as mean normalized counts ± SD from data reported in Kang *et al*, 2025 (Supp. Text ref. 4). (B) Cxt1-HA and Cxt2-FLAG band intensity normalized to actin. The quantitation shown is representative of two experiments, each with two independent strains.

### COMPLETE METHODS

#### Strain construction and growth

Strains are detailed in Table S2. All strains used in this study are mating type alpha. The *cxt1*Δ *cxt2*Δ deletion strains were made by sequential split-marker biolistic transformation (5) in JEC21 and KN99 (*CXT1* gene IDs: CNL04750, CKF44_05148; *CXT2* gene IDs: CND04030, CKF44_01283). *cxt1*Δ *cxt2*Δ was then backcrossed to the wild-type parent to obtain isogenic wild-type, *cxt1*Δ, *cxt2*Δ, and *cxt1*Δ *cxt2*Δ strains. For epitope-tagged strains, Cxt1 was C-terminally tagged with Myc by biolistic gene delivery (6) or with 6x-HA (as in pFSC6xHA) by CRISPR with short homology-directed repair (7). Cxt2-HA (pMSC-018) was episomally expressed under the actin promoter after electroporation (8, 9). Cxt2 was also C-terminally tagged with GSGS-2xFLAG in its endogenous locus with a drug resistance cassette in the Safe Haven 2 region by CRISPR as in ref. (10). Transformants were validated by drug selection and PCR amplification or whole genome sequencing.

Strains were grown on yeast extract-peptone-dextrose (YPD) plates for 2 days at 30°C. For liquid cultures, single colonies were inoculated into YPD and grown overnight at 30°C, with shaking. For capsule induction in host-like conditions, overnight YPD cultures were washed in PBS and grown in high-glucose DMEM (Sigma, D6429) at $10^6$ cells/mL for 24 h at 37°C and 5% $CO_2$, without shaking.

| Strain Description | Background | Strain ID |
|---|---|---|
| *ade2*Δ | JEC21 | TDY399 (JEC50) |
| WT | JEC21 | TDY2149 |
| *cxt1*Δ::*NAT* | JEC21 | TDY2147 |
| *cxt2*Δ::*G418* | JEC21 | TDY2148 |
| *cxt1*Δ::*NAT cxt2*Δ::*G418* | JEC21 | TDY2146 |
| WT | KN99 | TDY1025 |
| *cxt1*Δ::*NAT* | KN99 | TDY1023 |
| *cxt2*Δ::*G418* | KN99 | TDY1024 |
| *cxt1*Δ::*NAT cxt2*Δ::*G418* | KN99 | TDY1022 |
| *ade2*Δ *ura5*Δ *uxs1*Δ::*ADE2* | JEC21 | TDY532 (NE178) |
| *CXT1* replaced with $P_{CXT1}$-*CXT1-Myc G418* | JEC21 | TDY1299 |
| *TDY1299* + episomal $P_{ACT1}$-*CXT2-HA NAT* | JEC21 | TDY1300 |
| *CXT1*::*CXT1*-$HA_6$ *G418 CXT2*::*CXT2*-$FLAG_2$ *NAT* | KN99 | TDY3784 |

**Supplementary Table 2.** Strains used in this study. Drug resistance markers were *NAT* (nourseothricin acetyl transferase) or *G418* (neomycin resistance gene from Tn5); P, promoter of the gene indicated in the subscript; sequences encoding epitope tags were Myc, HA, or FLAG with the number of repeats indicated as subscripts.

**RNA interference**

RNA interference was performed as we previously described (1, 12, 13). Briefly, the cDNA for each RNAi target was amplified from JEC50 (12) and inserted into the pIBB103 vector (Genbank accession number HQ455038), which contains a G418 resistance cassette. The cloning site and *URA5* fragment are flanked by convergent, galactose-inducible *GAL7* promoters which drive bidirectional transcription, producing dsRNA that silences *URA5* and the target gene. 5-fluoroorotic acid (5-FOA) and G418 selection were used for RNAi selection and plasmid maintenance, respectively.

**Protein sequence alignment and structure analysis**

Cxt1 and Cxt2 protein sequences from JEC21 (CNL04750 and CND04030, respectively) and KN99 (CKF44_05148 and CKF44_01283, respectively) were obtained from FungiDB (2) and aligned using Needleman-Wunsch global alignment. Glycosyltransferase domains are marked with InterProScan and SMART predictions according to FungiDB annotations.

AlphaFold3 predicted structures (14, 15) were aligned using Pymol. The Template Modeling (TM) score (16) was determined with AlphaFold3 predicted structures using the TM-score online tool (https://zhanggroup.org/TM-score/).

**Phenotyping**

Overnight YPD-grown cultures were adjusted to $10^7$ cells/mL in PBS. 3 µL aliquots of 10-fold serial dilutions were spotted on YPD agar without or with 1.2 M NaCl, 0.01% SDS, 2.2 mM calcofluor white, or 10% DMSO (calcofluor white vehicle control). Plates were incubated at 30°C or 37°C for 72 h.

**Glycolipid preparation and analysis**

Glycosphingolipids were prepared and analyzed as we previously described (17). Briefly, cells were cultured for 3 days in YPD, washed once in cold water and twice in cold 20 mM sodium azide, and frozen. Cell pellets (50-70 g wet weight) were homogenized in 6 volumes of 1:1 (vol/vol) chloroform:methanol and the extracts evaporated under nitrogen. Glycosphingolipids were enriched, recovered in acidic fractions by ion-exchange chromatography, and purified by high-performance liquid chromatography (HPLC) as in references (18, 19). Crude acidic fractions were analyzed by high-performance TLC (HPTLC) on Silica Gel-60 TLC plates developed in 60:40:9 (vol/vol/vol) chloroform:methanol:water containing 0.002% (wt/vol) $CaCl_2$. Hexose-containing components were visualized by Bial's orcinol reagent.

**Glycoprotein preparation, oligosaccharide release, and mass spectrometry**

Cell lysates were prepared as we previously described (20), with all steps performed at 4°C with ice-cold buffer. Dense YPD cultures were sedimented and resuspended in 100 mM pH 8.0 Tris, 0.1 mM EDTA, and then disrupted by bead-beating. Samples were centrifuged (2,000 g; 25 m) to remove unbroken cells and debris, and the crude lysate (supernatant) was solubilized with 1% CHAPS (final concentration) by rotating for 2 h at 4°C. Insoluble material was removed by centrifugation (75,000 g; 45 min) and the extracts were dialyzed against 2 L ice-cold 50 mM ammonium bicarbonate buffer using 6-8 kDa Fisher brand regenerated cellulose dialysis tubing for 48 h (with buffer changes every 12 h). Dialyzed samples were lyophilized, then washed with ice-cold 80% acetone to remove residual detergent and other contaminants.

*O*-linked oligosaccharides were released by reductive β-elimination as and permethylated as we previously described (21). Lyophilized cell extract material was resuspended in 100 mM NaOH and 1 M sodium borohydride and incubated for 18 h at 45°C. The reaction was neutralized with 10% acetic acid on ice and desalted using an AG 50W-X8 cation-exchange resin. To remove borate, 10% acetic acid in methanol was added, then dried under nitrogen; the borate removal step was repeated for 4 additional rounds. The sample was then resuspended in 5% acetic acid and loaded onto a C18 cartridge column

(Mallinckrodt Baker). Flow-through and washes with 5% acetic acid were collected, combined, and dried. *N*-linked oligosaccharides were released from lyophilized cell extract material by PNGase F digestion and permethylated as we previously described (22, 23).

For Orbitrap MS analysis, MS2 spectra were acquired using total ion mapping (TIM) and full MS (MS1) spectra were collected at a resolution of 30,000. MS1 data were deconvoluted using Xtract software (Thermo Scientific) for quantification, and MS2 data from TIM were used for structural identification.

Nanospray ionization mass spectrometry (NSI-$MS^n$) was performed as previously described (20, 21). Permethylated glycan samples were dissolved in 1 mM NaOH in 50% MeOH and infused into a linear ion trap mass spectrometer equipped with an Orbitrap FT analyzer (LTQ-Orbitrap Discovery, ThermoFisher).

Mass spectra for these studies are available in GlycoPOST under accession number: GPST000698 (24). Glycomics data and metadata were obtained and are presented in accordance with MIRAGE standards and the Athens Guidelines (25, 26). Symbol and text nomenclature for representation of glycan structures is according to the Symbol Nomenclature for Glycans (SNFG) (27). The explicit identities of individual monosaccharide residues are assigned based on known fungal biosynthetic pathways. Data annotation and assignment of glycan accession identifiers was facilitated by the GRITS Toolbox (28), GlyTouCan (29), and GlyGen (30, 31).

**Capsule analysis**

Cell cultures were induced as described above, then washed, resuspended in PBS, and mixed with India ink (Higgins) in PBS (2:1 vol/vol) to visualize capsule by negative staining. Randomly chosen fields of cells were imaged on a ZEISS Axio Imager M2 fluorescence microscope. The areas of each cell body with and without capsule were measured by ImageJ; their radii were calculated for at least 80 cells per sample and plotted as a ratio.

**Transmission electron microscopy**

Ultrastructural analyses were performed as in reference (32). Strains were cultured overnight in YPD at 30°C, sedimented, and resuspended in in 2% glutaraldehyde (Polysciences) in 100 mM phosphate buffer, pH 7.2 for 1 h at room temperature and then overnight at 4°C. Samples were washed in phosphate buffer and postfixed in 1% osmium tetroxide (Polysciences Inc.) for 1 h. Samples were then rinsed extensively in $dH_20$, dehydrated in increasing concentrations of ethanol in water, and substituted in propylene oxide (PO) before being embedded in Eponate 12 resin. Sections of 90 nm were cut using a Leica Ultracut UCT ultramicrotome (Leica Microsystems), stained with uranyl acetate and lead citrate, and viewed with a JEOL 1200 EX transmission electron microscope (JEOL USA) equipped with an AMT 8-megapixel digital camera (Advanced Microscopy Techniques).

**GXM immunoblot**

To obtain shed GXM from conditioned media, cultures were grown in YPD at 30°C for 5 days. The conditioned media (supernatant) was separated from cell pellets by centrifugation at 3,000 rcf for 5 min. Conditioned media was then filtered through 0.22 μm syringe filters and dialyzed using MWCO 3kDa membranes for 5 days in MilliQ water at 4°C, changing the water twice daily.

GXM electrophoresis and immunoblot were performed as previously described (34). Dialyzed conditioned media was mixed 5:1 (v:v) with 6x DNA loading dye and samples were loaded in a 0.6% megabase agarose gel in 0.5x TBE (44.5 mM Tris base, 44.5 mM boric acid, 1 mM EDTA), separated by electrophoresis at 20 V for 16 h, and transferred to a positively charged nylon membrane overnight in 20x SSC (3 M NaCl, 0.3 M $Na_3$ citrate · $H_2O$, pH 7.0). Membranes were blocked with 5% milk in TBS for 1 h, immunoblotted with 1 μg/mL anti-GXM antibody 18B7 for 1 h, washed with TBS Tween-20, incubated with 1:10,000 680RD goat anti-mouse IgG (LI-COR IRDye, 926-68070) for 1 h, washed with TBS Tween-20, and visualized on a LI-COR 92120 Odyssey Infrared imaging system.

**Virulence studies**

All animal protocols were approved by the Washington University Institutional Animal Care and Use Committee, with care taken to minimize animal handling and discomfort. Groups of three or five 6-8-week-old female C57BL/6 mice from Jackson Laboratory were anesthetized by subcutaneous injection of 120 µL of 10mg/mL ketamine and 2 mg/mL xylazine in sterile PBS and intranasally infected with either 2.0 x $10^4$ cells (JEC21 background) or 2.5 x $10^4$ cells (KN99 background) in 50 µL PBS. Mice were monitored with care and humanely sacrificed 12 days post-infection; lungs and brains were harvested, homogenized, and plated on YPD agar to assess fungal organ burden. Colony-forming units (CFU) were counted, normalized to organ weight, and analyzed by ordinary one-way ANOVA with Tukey's multiple comparisons test.

**Immunofluorescence**

Immunofluorescence was performed as in reference (20). Cells were cultured overnight in minimal medium (0.17% w/v YNB without amino acids and $(NH_4)_2SO_4$, 0.5% w/v $(NH_4)_2SO_4$, 2% w/v glucose) at 30°C and fixed with 3.7% formaldehyde for 35 min at 30°C. 4 x $10^8$ cells were sedimented, washed with PBS, and incubated in fresh 3.7% formaldehyde for 35 min at room temperature. They were then washed, resuspended in Lysing Buffer (50 mM sodium citrate, pH 6.0, 1 M D-sorbitol, 2.5 µL/mL β-mercaptoethanol), and treated with 30 mg/mL Lysing Enzymes from *Trichoderma harzianum* (Sigma L-1412) for 1 h at 30°C. After digestion, cells were washed and resuspended in HS buffer (100 mM HEPES pH 7.0, 1 M D-sorbitol).

Washed cell suspensions were spotted in 20 μL aliquots onto 0.1% poly-L-lysine coated glass slides and incubated at room temperature for 30 min. In all subsequent steps, treatments were performed in 20 μL volumes and washes were by aspiration. Cells were permeabilized in HS buffer + 1% Triton X-100 for 10 min, washed, and blocked in 5% goat serum with 0.02% Tween-20 in PBS for 1 h before treatment with primary antibody ((1:300 α-Myc-Tag rabbit mAb IgG clone 71D10 (Cell Signaling #2278) or 1:100 or 300 α-HA mAb rat High Affinity (Roche #11867423001)) overnight at 4°C in a moist chamber. Afterwards, cells were washed with blocking buffer and stained with 1:1000 secondary antibody (α-Rabbit goat IgG AlexaFluor-488 (Invitrogen #A11008), α-Rabbit chicken IgG AlexaFluor-488 (Invitrogen #A21441), or α-Rat goat IgG AlexaFluor-594 (Invitrogen #A11007)) for 1.5 h. Following staining, cells were washed with blocking buffer, stained with 5 μg/mL DAPI (Sigma, D-8417) for 30 min, washed again, and treated with 4.5 μL Prolong Gold (Invitrogen, P-36930) for 24 h in the dark. Bright field and fluorescence images were acquired on a Zeiss Axioskop MOT Plus equipped with an AxioCam MRM 2.0 Hi-Res digital camera and AxioVision software.

**Protein extraction and immunoblotting**

YPD-grown cells were transferred to high-glucose DMEM at 37°C and 5% $CO_2$ and grown for 0, 30, 90, 180, or 1440 min before protein extraction. Cells were sedimented, washed in cold MilliQ water, and resuspended in ice-cold immunoprecipitation lysis buffer (100 mM Tris pH 7.6, 150 mM NaCl, 1 mM EDTA, 1% NP-40, and protease inhibitor cocktail (Sigma P8215)). Cells were next disrupted by bead-beating (8 rounds of 1 min bead beating followed by 1 min on ice) and unbroken cells were removed by centrifugation (500 g; 20 min; 4°C). The protein concentration of the resulting supernatant (cleared cell lysates) was determined by BCA (Pierce BCA Protein Assay Kit, 23225).

Protein samples were mixed with 5x SDS sample buffer containing beta-mercaptoethanol and boiled for 5 min. Samples were resolved on 4-15% SDS-PAGE gels and transferred to nitrocellulose membranes, which were blocked with 10% milk in TBS for 1 h, washed with TBS Tween-20, and incubated with primary antibodies (1:5,000 anti-HA antibody (Abcam, ab9110), 1:500 anti-FLAG (Millipore Sigma, F3165), and 1:1,000 anti-actin antibody (ThermoFisher, MA1-744)) overnight at 4°C. Membranes were washed with TBS Tween-20, incubated with secondary antibody (1:10,000 680RD goat anti-mouse IgG (LI-COR IRDye, 926-68070) or 1:10,000 800CW goat anti-rabbit IgG (LI-COR IRDye, 926-32211)) for 1 h at room temperature, washed with TBS, and visualized on a LI-COR 92120 Odyssey Infrared imaging system.

**Co-immunoprecipitation**

Cells were grown in YPD overnight (YPD-grown), transferred to host-like conditions for 30 min (DMEM-grown), and protein extraction was performed as described above. 500 μg of cleared cell lysates were incubated with 30 μL of washed Protein G agarose beads (Cell Signaling Technology, 37478), 1.5

µg (1.5 µL) anti-HA antibody (Abcam, ab9110), and fresh protease inhibitor cocktail for 1 h at room temperature. The beads were sedimented (14,000 g; 1 min) and washed three times with ice-cold wash buffer (0.05% TBS Tween-20). The unbound and first wash fractions were stored for further analysis. To elute bound proteins, beads were incubated with 100 µL elution buffer (2x SDS non-reducing sample buffer) for 10 m at 65°C. Eluates were separated from beads by centrifugation (14,000 g; 1 m) and treated with 2.5 µL beta-mercaptoethanol. Beads were then resuspended in 100 µL 2x SDS reducing sample buffer. All samples were boiled for 5 m before SDS-PAGE and transfer to nitrocellulose. Immunoblotting was performed as described above, with the primary antibodies anti-HA (Abcam, ab9110; 1:5,000 dilution) and anti-FLAG (Millipore Sigma, F3165; 1:2,000 dilution).

**Statistical methods**

Statistical analysis was performed using Prism (GraphPad) for ordinary one-way ANOVA with Tukey's multiple comparisons test.
